# Spatial Heterogeneity of Macrophages in the Human Lung

**DOI:** 10.1101/2025.05.30.657106

**Authors:** Patrick S. Hume, Katelyn H. Lyn-Kew, Elizabeth A. Wynn, Benjamin Steinhart, Jennifer Driscoll, Sean Jacobson, Peter M. Henson, Kara J. Mould, Camille M. Moore, William J. Janssen

## Abstract

**Rationale:** Transcriptionally-defined populations of interstitial macrophages (IMs) and airspace macrophages (AMs) have recently been identified in the human lung. However, the anatomic locations occupied by these populations (i.e. alveoli, pleura, airways, or arteries) have not been fully defined.

**Objectives:** To determine the distribution of transcriptionally-defined human macrophages in the major anatomical lung structures and to identify alterations in their distribution and programming induced by cigarette smoking.

**Methods:** Single-cell RNA sequencing was performed on lung tissue from eight human donors without pulmonary disease (four smokers and four nonsmokers). Microdissection was used to isolate distinct pulmonary anatomical structures from each lung: alveoli, pleura, airways, and arteries. Transcriptional profiles of subpopulations of interstitial macrophages (IMs) and alveolar macrophages (AMs) were analyzed based on their anatomical structure of origin and smoking status.

**Measurements and Main Results:** Five major IM and five AM subpopulations in human lungs are identified. We demonstrate significant differences in the accumulation patterns of each macrophage subset within anatomical structures, though each subset was detected in each. Immunofluorescent microscopy confirmed anatomical structure-specific accumulation patterns of IMs.

**Conclusions:** In this study, we highlight key differences in the accumulation of lung macrophage subpopulations in anatomical structures but find programming within macrophage subpopulations is largely conserved, regardless of structure of origin or smoking status. We also detect populations of inflammatory AMs and IMs which accumulate within the airways, but not the alveolar parenchyma, of human cigarette smokers. We introduce a novel three-tiered hierarchy nomenclature to distinguish transcriptionally defined human lung IM subsets as 1°) Monocyte-like vs Antigen Presenting, 2°) Quiescent vs Inflammatory, and 3°) FOLR2^high^ vs FOLR2^low^. This study is the first to report the fractional accumulation of human lung macrophage subsets by lung anatomical structure.

**Summary:** Lung anatomical structure-specific single cell RNA sequencing is introduced to identify and determine the local composition of human lung leukocytes, including 5 populations of human interstitial macrophages.

## Introduction

The human lungs are populated by a diverse array of leukocytes, including billions of macrophages (1) that are broadly defined based on their residence in the interstitium (interstitial macrophages, IMs) or airspaces (airspace or alveolar macrophages, AMs). These macrophages play key roles in maintaining homeostasis, promoting host defense, and coordinating inflammatory responses (2–5). They are also implicated in the pathogenesis of a myriad of lung diseases including COPD (6), asthma (7), pulmonary fibrosis, (8) and pulmonary hypertension (9). The remarkable diversity of macrophage functions is likely driven by a division of labor among cells occupying distinct lung structures and by the specialization of discrete macrophage subpopulations.

Using stereology, we previously quantified an abundance of human IMs in each of the major anatomical tissues (i.e. alveolar septa, pleura, airways, vessels) and AMs in the alveolar and airway lumens (1). However, this study was not designed to resolve specific IM subpopulations. We’ve since hypothesized that unique tissue niches drive macrophage identity and function in each lung structure (2, 10). This is supported by previous tissue histology approaches highlighting a subpopulation of IMs expressing Lyve1 residing near the vasculature (8). Our group has also reported a population of murine IMs expressing folate receptor β (Folr2) residing within the bronchovascular bundles (11). A limitation of imaging-based studies is that other macrophage subpopulations potentially coexist nearby but may be unidentified unless specifically targeted. Therefore, we set out to test whether unique macrophage subsets are enriched within specific anatomical structures of the human lung.

Recent investigations using the agnostic approach of single cell RNA sequencing (scRNA-seq) (8, 11–17) have confirmed the presence of multiple IM and AM subsets in both mice (8, 17, 18) and humans (8, 17, 19, 20). Within the interstitium, there appear to be two major subpopulations of IMs, with consensus emerging around expression of Timd4, Lyve-1 and Folr2 distinguishing “TLF” positive and “TLF” negative subsets (17). TLF-negative IMs can be further subdivided into at least two additional subsets: HLA^high^ and CCR2^high^, though the functions of these populations are less well defined (17, 19). Patterns of cytokine expression may be used to parse mouse IM populations even further (15). Along similar lines, several subpopulations of AMs have been identified in the healthy human lung, including homeostatic antigen presenting and inflammatory-primed populations, metallothionein-high and interferon-responsive cells (12, 13). Though instrumental to progressing the field, a key limitation of most recent transcriptomic investigations is their reliance upon either homogenized tissue that aggregates macrophages from all anatomical structures, or bronchoalveolar lavage that aggregates AMs located in both airway and alveolar lumens. Though not specifically interrogated for this purpose, a recent transcriptomic dataset suggests human macrophage composition varies by lung structure (21). No study to date has comprehensively evaluated relationships between transcriptionally defined macrophage subsets and pulmonary anatomical structure.

In this study, we report single-cell RNA sequencing of microdissected lung regions to map macrophage subsets across alveoli, pleura, airways, and arteries. We report a novel nomenclature to define a hierarchy of lung IM subsets, starting with two primary IM populations: Monocyte-like and Antigen Presenting. We identify distinct regional accumulation patterns for multiple macrophage subtypes and assess how smoking influences their distribution and transcriptional programming. These findings provide a spatial framework for human lung macrophage diversity and offer insights into how an environmental exposure shapes anatomical structure-specific immune landscapes.

## Methods

### Study Population

Studies used de-identified human lung tissue procured from organ donors that died from non-pulmonary causes, had clear chest x-rays and no history of pulmonary disease. Donors were classified as either nonsmokers or smokers. Smokers had ongoing, daily cigarette smoking for at least 10 pack-years, while nonsmokers had never smoked. Hematoxylin and eosin-stained sections from each lung were analyzed. Lungs with histologic evidence of acute or chronic disease, including emphysema, were excluded.

### Human Lung Tissue Isolation, Digestion and CD 45-Enrichment of Single Cells

Workflow is summarized in Figure 1A. The right upper lobe (RUL) was separated from the middle and lower lobes and visually inspected. The RUL bronchovascular bundle was then followed distally and carefully dissected from the surrounding alveolar tissue. The large airway was isolated and dissected from the adjacent pulmonary artery, which was also saved. To isolate distal alveolar tissue and pleura, a 3 cm^3^ piece of tissue was isolated from the distal lung. The pleural surface was carefully dissected and peeled away from the adjacent alveolar tissue and saved.

**Figure 1.**
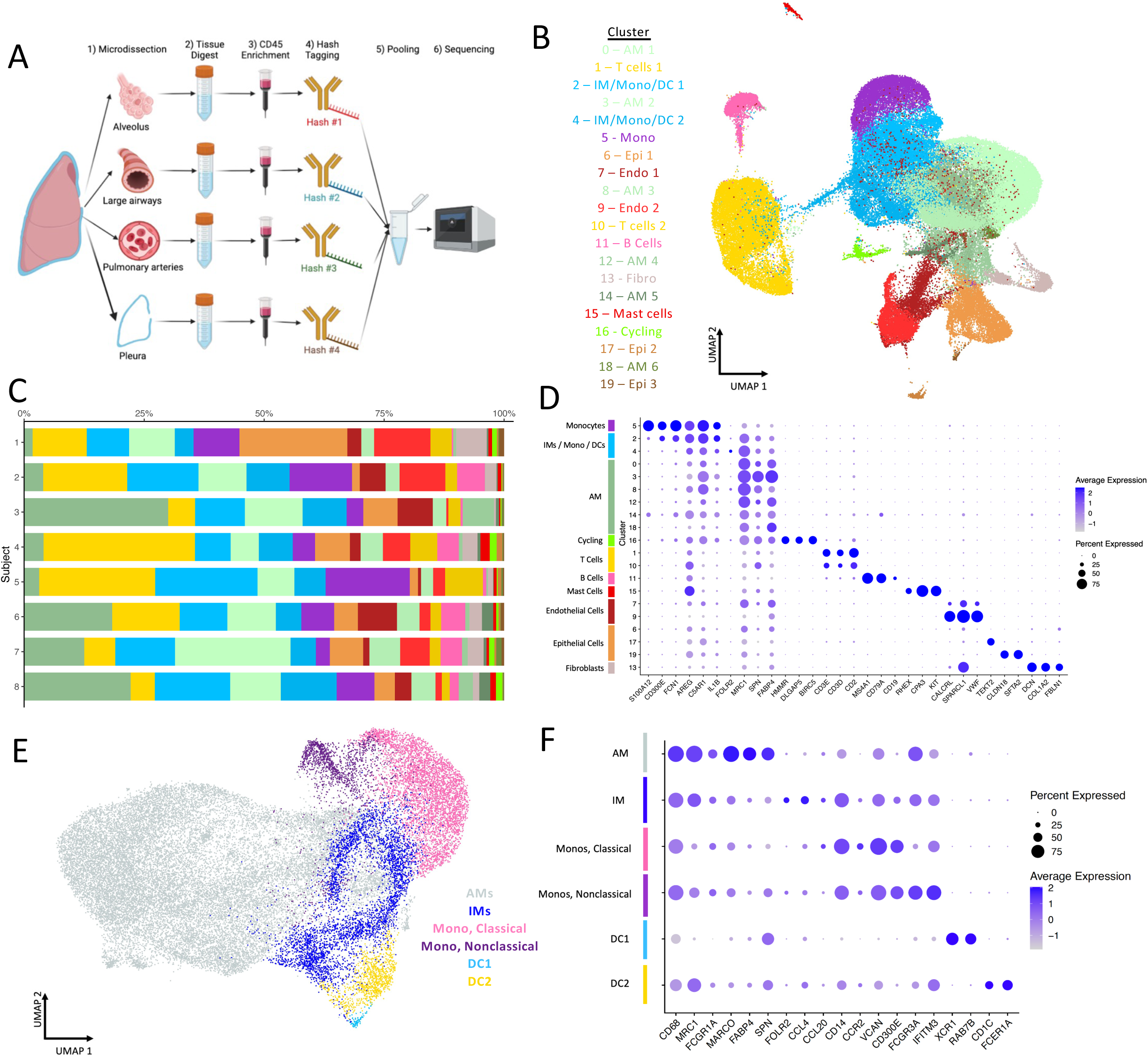
Single Cell RNA-seq of human lung tissue. A) Workflow to isolate lung leukocytes for single cell RNA sequencing. Right upper lobe from each study lung was isolated, then 1) microdissected it into 4 substructures: alveolus, large airways, pulmonary arteries, and pleura. 2) Tissue digestion yielded a single cell suspension. 3) CD45 enrichment was achieved with magnetic beads. 4) Cellular Hash-Tagging labeled cells from each structure with a structure-specific RNA oligo tag. Cells from each structure were pooled in equal fraction and sequenced. B) Uniform Manifold Approximation and Projection (UMAP) plot of clusters identified via the Seurat algorithm identifies 20 clusters of cells which were manually annotated and included populations interstitial macrophages (IMs), monocytes (Monos), dendritic cells (DCs) and airspace macrophages (AMs), epithelial cells (Epi), endothelial cells (Endo), T cells, B cells, fibroblasts (Fibro), mast cells, and cell with increased expression of cell cycle markers (Cycling), C) Fractional accumulation of each cluster in each of the 8 donors. D) Top differentially expressed marker genes for each of the major cell clusters. E) Myeloid populations identified via scRNA seq. with overlay showing subsets of: airspace macrophages (AMs), interstitial macrophages (IMs), classical monocytes (MonoC), nonclassical monocytes (MonoNC), dendritic cell 1 (DC1) and dendritic cell 2 (DC2). F) Characteristic differentially expressed marker genes for each myeloid population.

Individual anatomical structures were mechanically minced with scissors and a razor blade, as previously described (1). Minced tissue was transferred to gentleMACS C Tubes (130- 093-237, Miltenyi Biotec) and enzymatic digestion was achieved in DNAse I (0.2 mg/mL, #NC1539905, SigmaAldrich), Collagenase D (2.5 mg/mL, #11088882001, Sigma Aldrich) and 10 mL sterile phosphate buffered saline (PBS, #20-012-050, ThermoFisher) for 30 minutes at 37 C (1, 22, 23). GentleMACs columns were then loaded into the GentleMACS Dissociator (130-093- 235, Miltenyi Biotec) and homogenized via the manufacturer’s default lung protocol for 10 to 30s. Digested tissue was filtered through a sterile 100 μM filter (#CLS431752, Sigma Aldrich) by gravity and then centrifuged for 5 min at 300 x g, yielding a cell pellet.

CD45 enrichment was performed using MACS LS Columns (#130-042-401, Miltenyi Biotec). Briefly, cells were incubated with anti-CD45 microbeads (#130-045-801, Miltenyi Biotec) for 20 minutes at 4 C, then rinsed with chilled PBS. After preparing the MACS Columns and loading into the magnetic field, labeled cells were added to the column and filtered via gravity allowing cells labeled with anti-CD45 to be retained in the magnetic field. Three rinses with MACS buffer (0.5% bovine serum albumin [BSA, #A8577, Sigma], 2mM EDTA [#15575020, ThermoFisher] in PBS) were performed to remove unlabeled cells and then the columns were removed from the magnetic column for the elution of CD45-enriched cells.

### Flow Cytometry

Fluorescently conjugated antibodies recognizing human CD45, CD206, CD43 and CD169 were included to identify macrophages; CD1c to identify dendritic cells; and CD3, CD19, CD15 and NucBlue to exclude T cells, B cells, neutrophils and dead cells, respectively. See Table S1 for all antibody clone information. To quantify macrophages via flow cytometry, 10,000 Flashred microsphere beads (#FR06M, Bangs Laboratory) were suspended in solution and run with flow cytometry, as previously described. FACS data were acquired using an LSR Fortessa (BD Biosciences) and analyzed using FlowJo software (BD Biosciences).

### Thawing cells, Hash-Tagging and Submitting cells for Single Cell RNA Sequencing (scRNA-seq)

To batch multiple samples for single cell RNA sequencing, it was necessary to freeze cells following lung digestion and CD45 enrichment. This has previously been shown to have minimal effect on cellular programming (13). On the day of scRNA-seq capture, frozen alveolus, airway, artery and pleura samples from each human lung were thawed independently by immersion in 37C water bath for ∼3 min until ice was melted. Samples (1mL/sample) were transferred to a new 15 mL conical tube and 14 mL scRNA-seq buffer (PBS containing 0.04% BSA) was added slowly to prevent rapid osmotic shifts in the recently thawed cells. Cells were counted with hemocytometer and then 0.5 million cells from each sample were transferred to individual tubes for hash-tagging.

Hashing to label cells isolated from tissue subcomponents was achieved via the addition of antibodies targeting beta-2 microglobulin and CD298, which are expressed ubiquitously on mammalian leukocytes (24). Hash tagging antibodies were added to each sample to barcode major anatomical structure of origin. Alveoli samples received Hashtag 1 (#394601, BioLegend), airways received Hashtag 2 (#394603, BioLegend), arteries received Hashtag 3 (#394605, BioLegend) and pleura received Hashtag #4 (#394607, BioLegend). 5 μL TruStain FcX FC block (#422302, BioLegend) and 45 μL scRNA-seq buffer were added to each cell pellet for 10 min at 4C to block Fc receptors. Then, 1 μL of the appropriate Hashtag antibody and 149 μL scRNA-seq buffer was added to each sample, to bring the total volume to 150 μL. Samples were incubated for 30 min at 4C and then rinsed with 1mL scRNA-seq buffer per well. After centrifugation (300xg x 5 min at 4C), samples were resuspended in 50 μL staining buffer and counted with a hemocytometer. Each sample was diluted to 1000 cell / μL and then an equal volume of cells from each structure (alveolus, airways, arteries and pleura) were combined into one vial per lung donor sample which was submitted for scRNA-seq.

Flow cytometry was performed before CD45 enrichment, after CD45-enrichment and following freeze-thaw cycle to demonstrate leukocyte enrichment, conserved population distributions, and viability, respectively, throughout our workflow Figure S1A-C. Flow cytometry using fluorescently conjugated antibody analogs of the clones used for cell hashing demonstrated a consistent, high degree of cell labeling. Figure S1D.

Single cells were captured using a Chromium Box (10x Genomics) to a target of 18,000 cells / sample. The Chromium Single Cell 3’ NextGem V3.1 library and gel bead kit (#PN1000075, 10X Genomics) were used along with Bead Chip B (#PN-1000092, 10X Genomics) for cell capture and library construction. Libraries were sequenced with the Illumina NovaSEQ 6000 sequencer (Illumina) using an S4 flow cell (Illumina) and sequencing was conducted to a target depth of 50,000 reads per cell.

### Single Cell RNA Sequencing (scRNA-seq) Analysis: Pre-Processing and Quality Control

Initial pre-processing of the 8 samples’ scRNA-seq data, including demultiplexing, alignment to the hg38 human genome, UMI-based gene expression quantification, and counting of hashtags, was performed using Cell Ranger (version 5.0, 10x Genomics) (25). To ensure high quality single cells were used for downstream analysis, we filtered out cells with low numbers of genes (bottom 2% of each sample) or with greater than 40% of mapped reads originating from the mitochondrial genome. Although observations comprising multiple cells were removed during the cell selection stage, we additionally safeguarded against doublets by removing all cells with a UMI count greater than the 98th percentile of UMI counts for each sample. Prior to downstream analysis, select mitochondrial and ribosomal genes (genes beginning with MT-, MRPL, MRPS, RPL, or RPS) were removed. The final quality-controlled dataset consisted of 84,931 cells and 22,485 genes.

To account for differences in coverage across cells, we normalized and variance stabilized UMI counts for each subject using the SCTransform method in the Seurat R package (26). In addition to adjusting for sequencing depth, we also adjusted for the proportion of mitochondrial reads.

Cells were assigned to lung anatomical structure using the HTOdemux function in Seurat using k-medoid clustering on the normalized HTO values for initial clustering. For each hashtag oligo (HTO), we used the cluster with the lowest average value as the negative reference group and fit a negative binomial distribution to this cluster. We then used the 90^th^ percentiles of these distributions to call cells positive or negative for an HTO. Cells that were positive for more than one HTO were considered doublets. 10,298, 14,247, 11,390, and 7,943 cells were assigned to airway, alveoli, pleura, and arteries respectively. 36,670 cells were not classified and were excluded from lung structure-specific analyses.

scRNA-seq Analysis: Data integration, dimensionality reduction, clustering, and visualization Data from the 8 samples were combined using single cell integration implemented in Seurat version 4.0, which identifies mutual nearest neighbor (MNN) cells across pairwise subjects to use as “anchors” to perform batch correction. Integration was carried out using the top 30 dimensions from a canonical correlation analysis (CCA) based on SCTransform-normalized expression of the top 3,000 most informative genes, defined by gene dispersion using Seurat’s SelectIntegrationFeatures function. Integrated data were then clustered and visualized using the top 30 principal components (PCs). For visualization, we reduced variation to two dimensions using Uniform Manifold Approximation and Projection (UMAP; n.neighbors = 50, min.dist = 0.3). Unsupervised clustering was performed using a shared nearest neighbor (SNN) graph based on 30-nearest neighbors and then determining the number and composition of clusters using a smart local moving (SLM) algorithm (resolution = 0.4). This algorithm identified 20 clusters.

### scRNA-seq Analysis: Identification of cluster markers

To identify cluster markers, we carried out pairwise differential expression analysis comparing SCTransform-normalized expression in each cluster to all others using a Wilcoxon rank sum test. Markers were identified as genes exhibiting significant upregulation when compared against all other clusters, defined by having a Bonferroni adjusted p-value < 0.05 (adjusted for 22,485 genes), a log fold change > 0.25, and >10% of cells with detectable expression. Subject specific analyses were then performed to determine if the identified markers were consistent across subjects and to ensure that a single outlying subject did not drive marker results. Subject specific analyses were conducted by comparing SCTransform-normalized expression in each cluster to all others using a Wilcoxon rank sum test separately for each subject. Again, p-values were Bonferroni adjusted for 22,485 genes and the number of subjects with adjusted p-values < 0.05 for each marker was tabulated. Markers were considered “conserved” if they had adjusted p-values < 0.05 for at least 4 of 8 subjects. This analysis was performed using Seurat’s FindConservedMarkers function.

### scRNA-seq Analysis: Sub-clustering

We re-clustered subsets of cells to further characterize cell types that may have been obscured by heterogeneity among the broad cell types present in the full dataset. We sub-clustered myeloid cell populations (overall clusters: 0, 2, 3, 4, 5, 8, 12, 16, 18; N=33,884 cells). Clustering and marker finding were performed on these subsets using the same approach described above. Macrophages were clustered using 30 PCs with SLM resolution 0.4.

Further sub-clustering was performed to identify IM subsets. The clusters, PCs, and SLM resolution for the 3 iterations are as follows: 1) 30 PCs with SLM resolution 0.4 (IM-like populations from the macrophage sub-clustering: 2, 4, 6; N=11,417 cells); 2) 30 PCs with SLM resolution 0.8 (Clusters from 1^st^ IM sub-clustering: 1, 2, 3, 4, 6 to remove AM-like and DC clusters; N=7,669 cells) and 3) 30 PCs with SLM resolution 0.4 (clusters from 2^nd^ IM sub- clustering: 2, 3, 5, 6, 7, 8 to remove additional AM- and monocyte-like clusters, N=4,402 cells). One subcluster of low-quality cells was removed after the final iteration (cluster 5), leaving N=3,982 cells. Sub-cluster markers were identified as described above, comparing against all IMs and against all other cells in the full dataset.

In addition, we performed two iterations of sub-clustering of the airspace macrophages as follows: 1) 30 PCs with SLM resolution 0.15 (AM-like populations from myeloid sub-clustering [clusters 0, 1, 5, 9, 10, 11], 1^st^ iteration of IM sub-clustering [cluster 0] and 2^nd^ iteration of the

IM sub-clustering [clusters 0 and 1]; N= 20,833 cells) and 2) 30 PCs with SLM resolution 0.09 (clusters from 1^st^ AM sub-clustering: 0, 2, 3, 4, 5, 6 to remove low-quality cells; N=15,180 cells). Sub-cluster markers were identified as described above, comparing against all AMs.

### scRNA-seq Analysis: Label transfer to PBMC data

To determine if our IM sub-clusters were distinct from circulating monocytes, we performed label transfer analysis using Seurat to project cell type labels from our single cell dataset onto a publicly available PCMC scRNA-seq dataset (10X Genomics (27)). To identify transfer anchors, we found the top 2000 most variable features using the VST method and used 30 PCs for label transfer.

### scRNA-seq Analysis: Differential Expression between IM Subsets and Tissue Locations

To further distinguish between IM subclusters, we identified marker genes that differentiated each pair of IM subclusters, again using a Wilcoxon rank-sum test performed independently for each subject. Markers were considered conserved based on the same criteria previously described. We then clustered all unique marker genes identified from any pairwise comparison using Euclidean distance with Ward’s method (28) based on SCTransform-normalized expression data from all IM cells. A heatmap of the clustered genes revealed gene programs with similar transcriptional patterns, which were defined using cut points in the hierarchical dendrogram. For each program, pathway analysis was performed to infer associated cellular functions.

We applied the same subject-level differential expression testing and marker conservation criteria for additional comparisons including: 1) groups of IM subclusters defined by shared gene program expression (e.g., monocyte-like vs. antigen-presenting subsets); 2) pairwise comparisons of each tissue type within the IM population; and 3) comparisons of the monocyte-like and antigen-presenting IM subsets to monocyte and dendritic cells. For the third comparison, we additionally clustered the conserved marker genes and visualized them in a heatmap using the methods described above.

### scRNA-seq Analysis: Differential Expression between Smokers and Non-Smokers

We compared expression between smokers and non-smokers for each IM and AM subset using a pseudo-bulk approach to account for clustering of cells within subjects (29, 30). For each subset, we summed counts for each gene across all cells for a subject. We then removed lowly expressed genes that did not have counts per million > 1 in at least 4 samples and compared expression between smokers and non-smokers using negative binomial models implemented in edgeR. The Benjamini-Hochberg method (31) was used to control the false discovery rate.

### Pathway Enrichment Analysis

Pathway enrichment analysis of marker and DEG lists was performed using EnrichR with the Gene Ontology (GO) Biological Processes database 2023 (32).

### scRNA-seq Analysis: Comparison of macrophage proportions between lung structures

The fractional composition of each IM or AM subset per anatomical structure was determined for each of the 8 lung donors. This fractional composition was averaged across each of the 8 donors, resulting in the average fractional composition of each subset, per structure. Statistically significant differences in macrophage subset proportions, per structure, were detected via 2-way ANOVA with Bonferroni’s Multiple Comparisons Testing to determine P<.05.

### Tissue Histology & Multiplex Imaging

Formalin-fixed paraffin-embedded (FFPE) human lung tissue sections were cut into 5 μm-thick slices. Tissue was deparaffinized, rehydrated, and subjected to antigen retrieval using Dako Target Retrieval Solution, citrate pH 6.0 (Agilent Dako). Slides were stained with DAPI, tomato lectin, FolR2, CCL4, CD206, and CD43 using antibody clones detailed in Table S1. TrueVIEW Autofluorescence Quenching kit (Vector Laboratories) was used to reduce background fluorescence. Four-channel fluorescent images were captured using VS200 whole-slide scanning microscope (Olympus). Coverslips were then removed by shaking slides in TTBS and slides were washed to remove any remaining mounting media. 30 minutes of a 10 mg/mL LiBH4 in diH2O (Radtke) was applied to slides to bleach fluorescent molecules. Repeat imaging confirmed signal bleaching. Slides were then re-stained and re-imaged to obtain multiplex imaging.

Slide images were imported to image analysis software (Adobe Photoshop) for alignment and composite image formation and counting. Quantification of IMs was performed on 3 separate airways, arteries, pleura, and alveoli across 6 donor tissues without disease. Only cells with a visible nucleus and CD206 staining were considered macrophages. IMs were defined by CD206 positive and CD43 negative staining, then identified as Frβ+ or - and CCL4+ or -.

## Results

### Human Lung Tissue Selection and Structure-Specific Cell Labeling

Single cell RNA sequencing (scRNA-Seq) was performed on tissues from eight organ donors (Table 1). Four donors were never smokers, and four were current smokers with a mean cigarette smoke exposure of 28 (SD 15) pack-years, which did not vary significantly by sex. The average donor age was 56 (SD 7) years and did not vary significantly by sex or smoking status. Anatomical lung tissue structures (alveoli, pleura, airways and arteries) were individually dissected then mechanically and enzymatically digested to achieve single cell suspensions, Figure 1A. CD45+ leukocytes were enriched using magnetic bead columns and barcoded using a cell hashing antibody (hash tag oligo, HTO) unique to each structure. Finally, for each donor, samples from each of the four tissue structures were combined in equal fraction for scRNA-Seq.

**Table 1.** Demographics. Human lung tissue was donated from deceased male (M) or female (F) individuals without underlying disease who were either nonsmokers (NS) or chronic daily cigarette smokers (S). Pack years is defined as the number of packs of cigarettes smoked daily multiplied by the total number of years smoked.

| # | Age | Sex | Smoker | Pack years |
| --- | --- | --- | --- | --- |
| 1 | 50 | M | NS | 0 |
| 2 | 50 | M | NS | 0 |
| 3 | 61 | F | NS | 0 |
| 4 | 60 | M | NS | 0 |
| 5 | 59 | F | S | 12.5 |
| 6 | 60 | M | S | 20 |
| 7 | 42 | M | S | 36 |
| 8 | 61 | F | S | 45 |
| Total | - | - | - | - |
| Mean (Nonsmoker) | 55 | - | - | 0 |
| Mean (Smoker) | 56 | - | - | 28.4 |

### Single Cell RNA Sequencing of Leukocyte-Enriched Lung Tissue Identifies Tissue-Resident Myeloid Populations

Following single cell sequencing, data processing and quality control, a total of 84,931 cells were identified. Unsupervised clustering using Seurat (33) identified 20 distinct cell clusters (Figure 1B). Each cluster was represented in all 8 subjects, Figure 1C. Conserved markers for 10 major cell types were identified and used for annotation Figure 1D.

Sub-clustering was performed on cells of the myeloid lineage to resolve six major cell types (classical monocytes, nonclassical monocytes, AMs, IMs, DC1 and DC2), Figure 1E. Resultant sub-clusters were annotated based on conserved marker expression (Figure 1F) (34–37). Macrophages exhibited high expression of quintessential marker genes including MRC1, CD68, MARCO and FCGR1A (CD64). AMs were further distinguished from IMs by their high expression of FABP4 and SPN (13). Because we defined AM based on validated markers of tissue-resident AM, it is possible that small numbers of recruited AMs were misidentified as IM, as have been reported in the healthy human lung (13). To verify that the cells we defined as IMs were distinct from blood monocytes, we aligned cells from our dataset to a publicly available dataset of peripheral blood cells (27) using label transfer methods (33). As anticipated, cells that we identified as monocytes, T cells, B cells and NK cells in the lungs aligned with their expected cell populations in the peripheral blood dataset. Importantly, cells that we defined as IMs had no overlap with peripheral blood cells, supporting the assertion that these IMs were isolated from lung tissue and are not blood monocytes.

### Single Cell RNA Sequencing Detects 5 Distinct Interstitial Macrophage Clusters within a 3-teir Hierarchy

To characterize human lung IM heterogeneity, we subclustered the myeloid cells identified as IMs. Five distinct IM clusters were identified, Figure 2A. We analyzed the gene expression patterns of each cluster, incorporating genes previously associated with IM subsets (such as FOLR2, LYVE1, CCR2, HLA-DRB1)(17) and DEGs identified in our dataset as distinguishing IM clusters (i.e. VCAN, FCN1, CCL4, IL1B, C1QC), Figure 2B. Distinct transcriptional patterns emerged, for example clusters 1 and 2 expressed gene typically associated with monocytes (VCAN, CCR2, FCN1), clusters 3 and 4 expressed T-L-F markers (FOLR2 and LYVE1) and clusters 2 and 3 expressed inflammatory markers (CCL4, IL1B).

**Figure 2.**
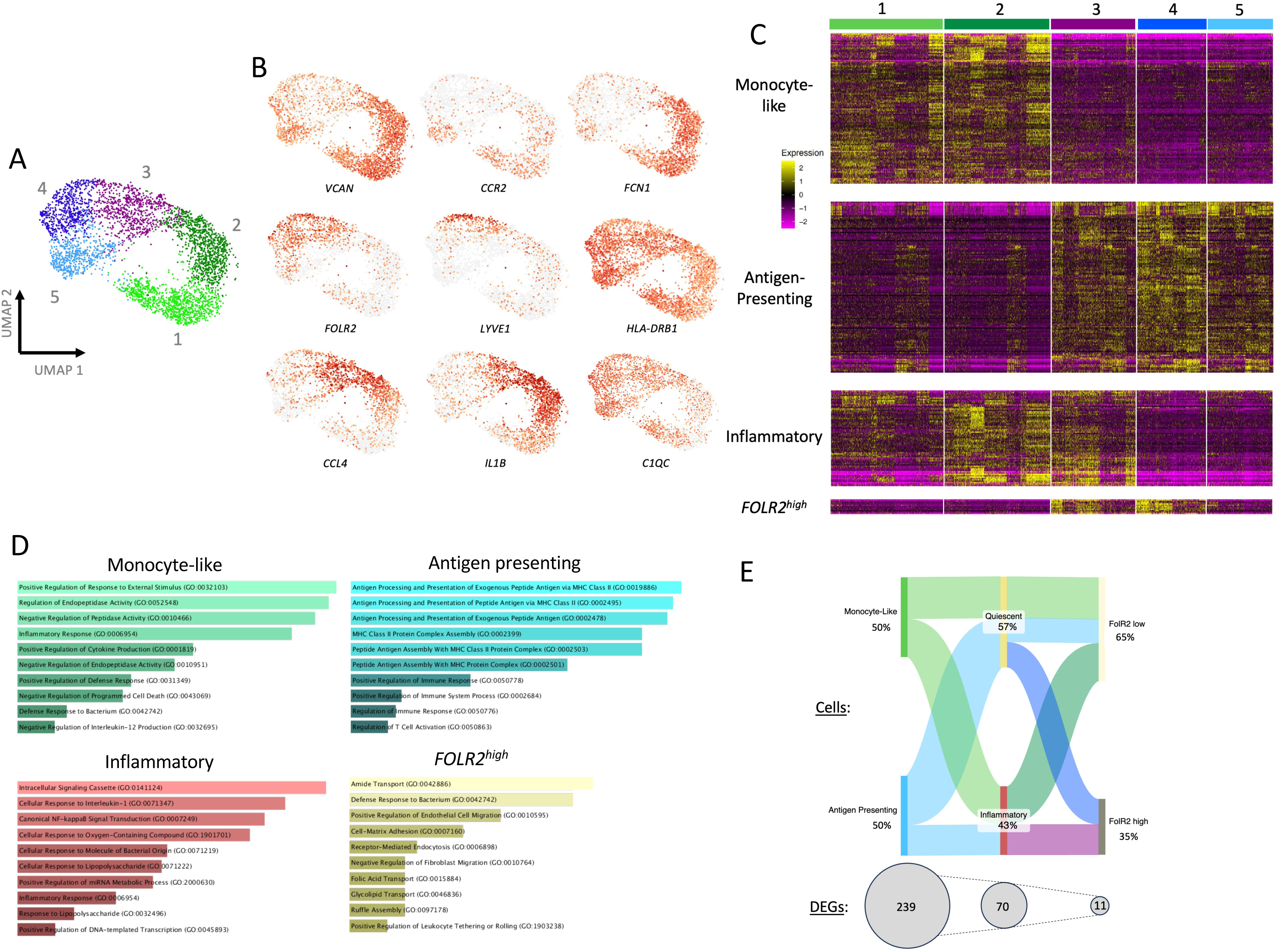
scRNA-seq identifies 5 Interstitial Macrophage (IM) populations in the human lung. A) UMAP projection of 5 IM clusters identified via Seurat algorithm. B) Selected gene features with variable expression amongst human lung IMs with single cells colored on a gradient from grey (no expression) to red (high expression) C) Heat map projection of all significant differentially expressed genes (DEGs) determined by pair-wise comparison of each IM subset. Four distinct programming signatures are identified: Monocyte-like, Inflammatory, Antigen presenting, and *FOLR2*^high^. D) Gene Ontology (GO) Pathways analysis performed on all statistically significant (P<0.05) DEGs defining each of the four core IM programming signature. E) Sankey plot illustrating (top) proportion of IM cells within each tiered nomenclature: 1°) monocyte-like vs antigen presenting, 2°) inflammatory vs quiescent and 3°) *FOLR2*^high^ vs *FOLR2*^low^ and (bottom) number of differentially expressed genes (DEGs) distinguishing each tier. Percentage represents the fractional cellular composition of each tier.

To clarify the transcriptional signatures defining IM cluster identity, we conducted all possible pairwise comparisons of gene expression profiles between IM clusters to identify differentially expressed genes (DEGs). Hierarchical clustering of the resulting 320 DEGs revealed four key transcriptional programs, Figure 2C. Pathway analysis of each program was performed to infer associated cellular functions, Figure 2D. Clusters 1 and 2 exhibited similar expression of ’Monocyte-like’ genes and upregulated pathways suggesting cytokine responsive and inflammatory activity. Clusters 3, 4, and 5 were enriched for genes associated with enhanced ’Antigen Presenting’ capacity and revealed several upregulated pathways suggesting antigen processing and presentation activity. Clusters 2 and 3 shared expression of ’Inflammatory’ cytokines such as CCL4 and IL1B and have increased pathways suggesting response to IL1 and lipopolysaccharide, NFκB activation and inflammatory signaling. Finally, Clusters 3 and 4 were distinguished by high expression of only 11 genes, including FOLR2, LYVE1, and SELENOP. Analyses of pathways for this programming signature is of questionable utility given the low number of DEGs. Thus, we have named this programming signature ‘FOLR2^high^’, rather than attempting to ascribe similarity to other cell types or predict function.

We sought to clarify the relative importance of each of these four transcriptional signatures in shaping IM identity. The greatest number of DEGs distinguished Monocyte-like and Antigen Presenting programming, with 114 and 125 genes respectively. This greatly exceeds the number of genes distinguishing Inflammatory IM programming (70 DEGs) or FOLR2^high^ programming (11 DEGs), Figure 2E. Notably, when Monocyte-like IMs (i.e., clusters 1 and 2) were aggregated and compared in a pairwise fashion with monocytes from our single-cell RNA-seq dataset, 190 DEGs were identified, fewer than 329 genes distinguishing Monocyte-like from Antigen Presenting IMs, Figure S2. Similarly, comparison of aggregated Antigen Presenting IMs (i.e., clusters 3–5) with dendritic cells yielded 216 DEGs, also fewer than the DEGs distinguishing the two IM subsets. These findings support a model in which two primary (1°) human lung IM populations exist: Monocyte-like IMs (clusters 1 and 2) and Antigen Presenting IMs (clusters 3– 5). The most significantly upregulated genes in Monocyte-like IMs include FCN1, VCAN, THBS1, S100A9, and S100A8. In contrast, Antigen Presenting IMs are enriched for C1QC, C1QA, C1QB, CD74, A2M, HLA-DPA1, HLA-DRA, and HLA-DQB1. Of note, macrophages expressing FOLR2 and LYVE1, consistent with previously reported TLF+ IMs (17), are localized within the Antigen Presenting IM subset, whereas CCR2+ IMs are found within the Monocyte-like IM subset. However, none of these markers (FOLR2, LYVE1, or CCR2) ranked among the top 30 DEGs distinguishing Antigen Presenting from Monocyte-like IMs.

Within each primary IM population (i.e., Monocyte-like or Antigen Presenting), we identified IM subclusters with and without the Inflammatory transcriptomic signature Figure 2C. For example, cluster 2 within Monocyte-like IMs and cluster 3 within Antigen Presenting IMs expresses the Inflammatory transcriptomic signature, while the other clusters do not. Because this Inflammatory signature is defined by upregulation of a set of 70 genes, far fewer than the 239 DEGs that drive Monocyte-like vs Antigen Presenting identity, we propose Inflammatory vs Quiescent programming as a secondary (2°) distinction downstream of Monocyte-like vs Antigen Presenting IMs, Figure 2E. Finally, the distinction between IMs with the FOLR2^high^ signature was driven by only 11 DEGs and was exclusive to a subset of Antigen Presenting IMs. Accordingly, we propose a tertiary (3°) level of IM identity: FOLR2^high^ versus FOLR2^low^ IMs, Figure 2E. In summary, we propose the following 3-teir hierarchy to define human lung IMs: 1° Monocyte-like vs Antigen Presenting, 2° Quiescent vs Inflammatory and 3° FOLR2^low^ versus FOLR2^high^.

### Location Mapping Identifies Spatial Heterogeneity of Interstitial Macrophages

To determine whether IM subsets localize to specific anatomic structures, we used structure-of-origin hashtags to assess the fractional composition of IM clusters within each region. As shown in Figure 4A, all five IM clusters were detected in each anatomical structure. However, the proportion of some IM subsets varied significantly by location. Quiescent Monocyte-like IMs (cluster 1) were the most enriched IM subset within the alveoli (30% of IMs, P<0.005) and pleura (37% of IMs, P<0.0001), Figure 4B. Inflammatory Monocyte-like IMs (cluster 2) predominated in the airways (29% of IMs) while Inflammatory Antigen Presenting IMs (cluster 3) where highly enriched in the arterial walls (47% of IMs, P<0.0001). Other IM subsets were distributed in roughly equal proportions across anatomical structures.

**Figure 4.**
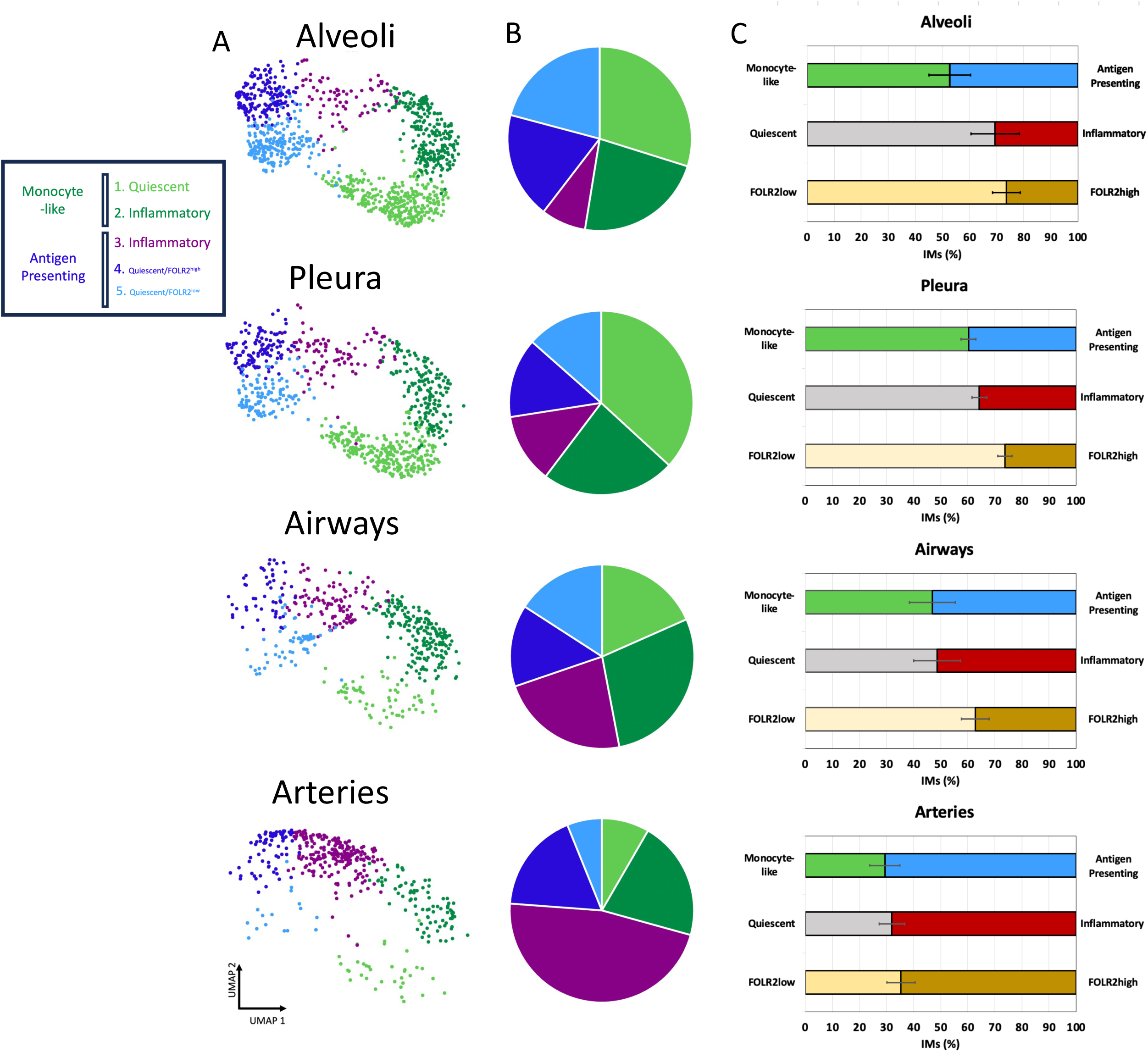
Spatial heterogeneity of human lung interstitial macrophage (IM) subsets by scRNA-seq. Column A) UMAP projections of each of the 5 IM clusters split by structure-of-origin hashtag as originating from Alveoli, Pleura, Airway walls or arterial wall. Column B) Pie charts expressing the fractional composition of each IM subset per structure. Key of colors denoting IM clusters in A) and B) is shown. Column C) Fraction of the number of IMs detected in each lung structure by hierarchical designation as Monocyte-like (green) vs Antigen Presenting (blue), Quiescent (grey) vs Inflammatory (red) and FOLR2^low^ (tan) vs FOLR2^high^ (brown).

Next, IM localization was cataloged by primary (1°), secondary (2°), and tertiary (3°) transcriptomic signatures, Figure 4C. In the alveoli, an equal proportion of Monocyte-like and Antigen Presenting IMs were detected but there was accumulation of Quiescent (69% of IMs) and FOLR2^low^ (74% of IMs). In the pleura, Monocyte-like IMs were slightly more abundant (60%) than Antigen Presenting IMs, with a similar accumulation pattern of Quiescent and FOLR^low^ IMs to the alveoli. In the airways, the fraction of FOLR2^low^ cells was modestly enriched (63%) but the fractions of Antigen Presenting vs Monocyte-like and Quiescent vs Inflammatory IMs were equal. The arterial wall showed a strong accumulation of Antigen Presenting (71%), Inflammatory (68%), and FOLR2^high^ (65%) IMs.

To determine whether IM programming varied by anatomical structure, we performed pairwise comparisons of transcriptional signatures between IMs that originated from different lung anatomical structures—for example, comparing IMs from the alveoli versus the airways. Nine significant DEGs were identified across all comparisons. Five genes (PLTP, RNASE1, CCL4, TNFAIP3, LIL4B5) were significantly upregulated in vascular IMs relative to those from alveoli, two genes (LYVE1, SELENOP), were upregulated in vascular IMs compared to both pleural and alveolar IMs, and two genes (LYZ, CAPG) were upregulated in alveolar IMs relative to vascular IMs.

CAPG appeared upregulated in Antigen Presenting IMs, and LYZ was upregulated in Monocyte-like IMs from the alveolar and pleural structures, but not from the airways or arteries, Figure S3. The inflammatory genes CCL4 and TFAIP3 were increased in Antigen Presenting IMs derived from the airways and vessels, but not from the alveoli or pleura. Five genes (LILRB5, LYVE1, PLTP, RNASE1, and SELENOP) were consistently upregulated in all IMs originating from the arterial structure. Overall, this relative paucity of DEGs suggests that programming within each IM subset is largely conserved, regardless of structure of origin.

Fluorescent Microscopy Confirms IM Subset Heterogeneity within Lung Anatomical Structures Immunofluorescent microscopy visualized IM subsets within human lung tissue sections to a) confirm the ability to detect IM subsets defined transcriptionally and b) corroborate the structural accumulation patterns predicted by scRNA-seq. As we have previously reported, positive staining for CD206 (macrophage mannose receptor, MMR) serves as a robust marker of all lung macrophages in histology sections (1). To identify protein targets to resolve IM subsets, we screened DEGs of each IM subset in our scRNA-seq dataset. Most Antigen Presenting IMs were distinguishable from Monocyte-like IMs by positive staining for folate receptor beta (FRβ), encoded by the FOLR2 gene. Inflammatory subsets of both Antigen Presenting and Monocyte-like IMs were resolved by positive tissue staining for CCL4. Thus, we developed the following strategy to detect IM subtypes in tissue sections: Inflammatory Monocyte-like IMs (cluster 2) as CD206+/FRβ-/CCL4+, Inflammatory FOLR^high^ Antigen Presenting IMs (cluster 3) are CD206+/FRβ+/CCL4+ and Quiescent FOLR^high^ Antigen Presenting IMs (cluster 4) are CD206+/FRβ+/CCL4-, Figure 5A-D. This staining strategy does not distinguish Quiescent Monocyte-like IMs (cluster 1) from Quiescent, FOLR2^low^ Antigen Presenting IMs (cluster 5), as both are CD206+/FRβ+/CCL4+. Cluster 5 lacked unique markers as its eight upregulated DEGs (e.g., HLA-DPA1, HLA-DPB1) were also expressed across many other IMs, while potential markers for cluster 1 (e.g., VCAN, AREG, EREG, CD300E) lacked sufficient specificity. As a result, neither population could be uniquely identified by positive staining. So, we herein report an indeterminant IM cluster detected as CD206+/FRβ-/CCL4- in histology studies which we predict from scRNA-seq to represent a combined clusters 1 and 5. scRNA-seq predicts the ratio of IM subsets within this indeterminant population would be 63% cluster 1 (Monocyte-like) : 37% cluster 5 (Antigen Presenting). Aside from this limitation, our staining strategy is adequate to distinguish Quiescent from Inflammatory IMs and FOLR2^low^ from FOLR2^high^ IMs.

**Figure 5.**
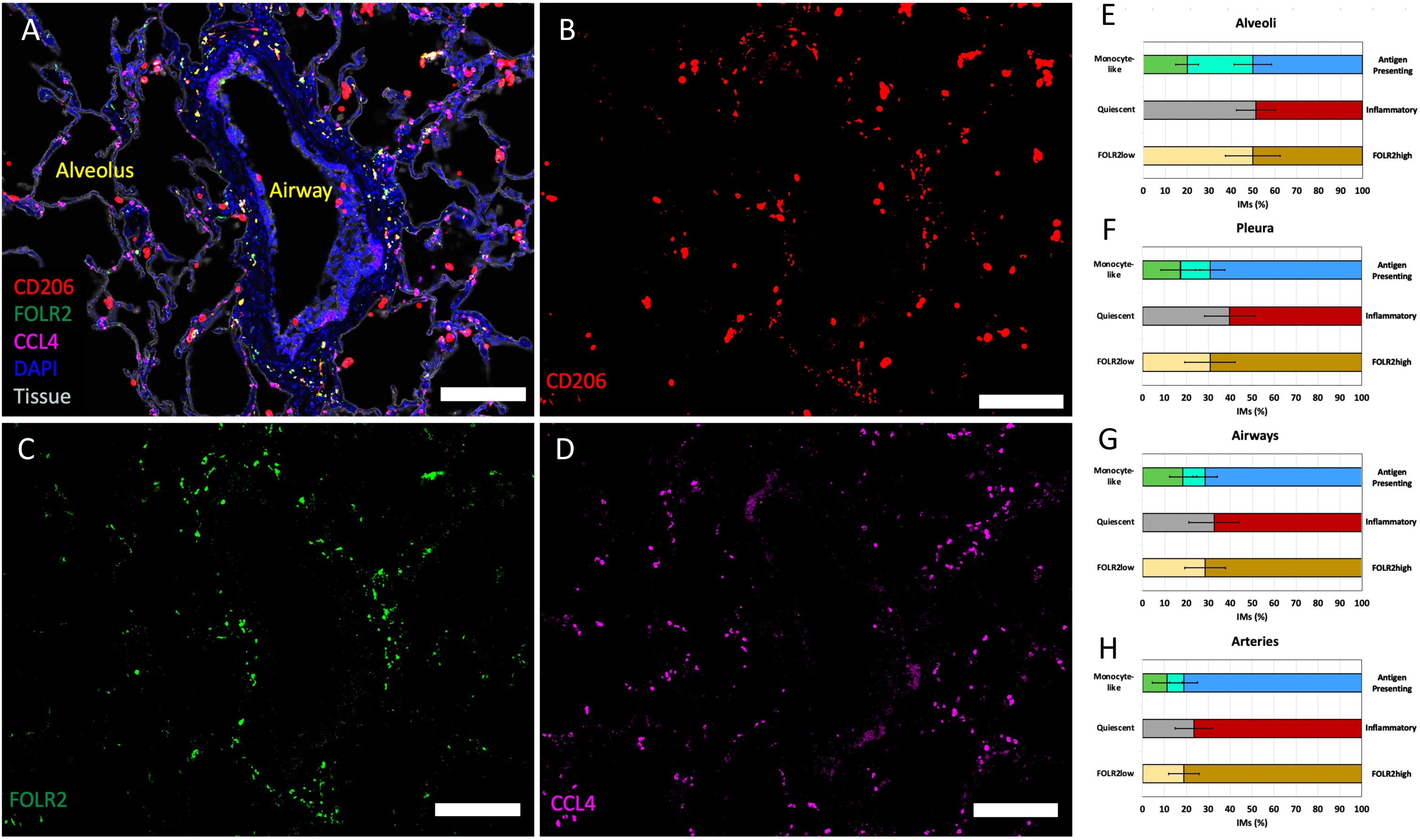
Immunofluorescent microscopy visualizes IM subsets in human lung tissue layers. A-D) Mosaic of immunofluorescent images of a representative view of stained human lung tissue showing an airway and alveoli. Scale = 200μm. A) View of all merged fluorescent channels [DAPI (blue), tissue lectin (white), CD206 (red), FOLR2 (green) and CCL4 (purple. Followed by single channel views showing B) CD206 (red), C) FOLR2 (green), D) CCL4 (purple). E-H) Fraction of IMs detected in each lung structure by hierarchical designation as Monocyte-like (green) vs Antigen Presenting (blue), Quiescent (grey) vs Inflammatory (red) and FOLR2^low^ (tan) vs FOLR2^high^ (brown) in E) Alveoli, F) Pleura, G) Airways and H) Arteries. Bright green bar between Monocyte-like vs Antigen Presenting fractions represents an “Indeterminant” population of cells unable to be distinguished by our staining panel.

Like scRNA-seq, microscopy detected each IM subtype in every major lung anatomical structure and revealed variable accumulation patterns by location, Figure 5E–H. In the alveolus, an approximately equal distribution of Quiescent vs. Inflammatory and FOLR2^low^ vs. FOLR2^high^ IMs was observed, Figure 5E. Microscopy revealed a higher proportion of Antigen Presenting IMs in the pleura (70% by microscopy vs. 40% by scRNA-seq, Figure 5F) and airways (71% vs. 53%, Figure 5G) than predicted by scRNA-seq. Within the vessels, IMs were predominantly Antigen Presenting, Inflammatory, and FOLR2^high^, Figure 5H.

### Chronic Cigarette Smoking Results in Subtle Alterations to IM Subset Distribution and Programming

To test the hypothesis that smoking alters the distribution and programming of macrophage subsets in human lungs, our experiment included specimens from 4 daily cigarette smokers and 4 nonsmokers. Initial analyses performed independently of structure-of-origin found the relative abundance of each of the 5 IM clusters did not vary significantly between smokers and nonsmokers, Figure 6A. Next, the transcriptional profiles of cells within each IM cluster were compared pairwise based on smoking status. Amongst Monocyte-like IMs (clusters 1 & 2), only 9 out of the more than 12,000 genes analyzed were significantly altered by smoking status, Figure S4. Amongst the subsets of Antigen Presenting IMs, a total of 29 DEGs distinguished smokers vs nonsmokers within either cluster 3 (Figure 6B), cluster 4 (Figure 6C) or cluster 5 (Figure 6D). In each of these three Antigen Presenting IM clusters, IFI27 and HLA-DQB2 were decreased in smokers. In cluster 4 (Quiescent FOLR2^high^ Antigen Presenting IMs), CCL18, CCL13 and several other genes were reduced in smokers, Figure 6C. PRR4, IFI27 and CX3CR1 were amongst the most significantly downregulated genes in cluster 5 (Quiescent FOLR^low^ Antigen Presenting IMs), Figure 6D.

**Figure 6.**
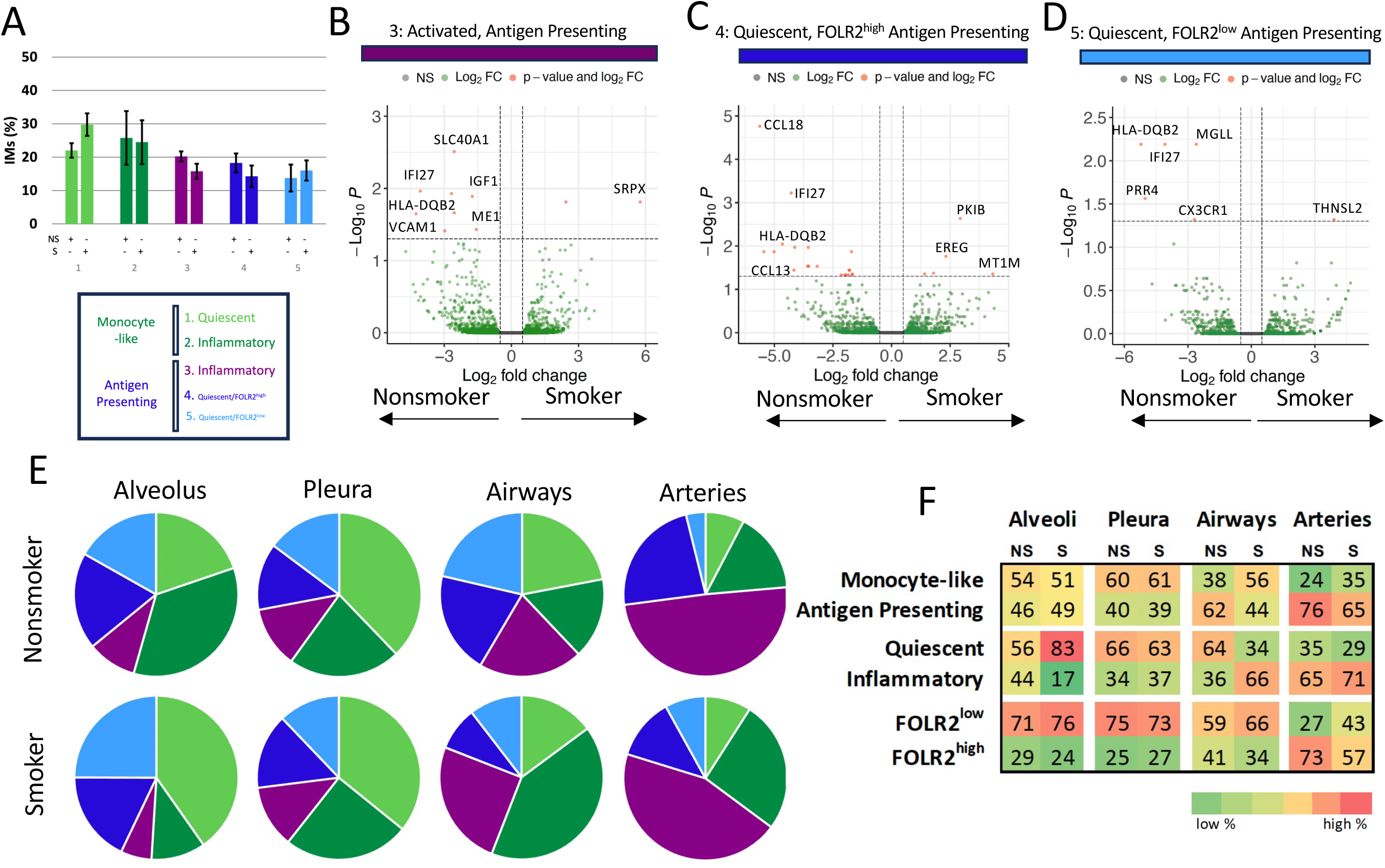
Influence of smoking status on human interstitial macrophage (IM) populations A) Percentage of each of the five interstitial macrophage (IM) clusters aggregated from all locations, split by donor status as Nonsmoker (top) or Smoker (bottom). B-D) Volcano plots highlighting all genes expressed by subsets of antigen presenting IMs compared pairwise between nonsmoker and smoker donors. X-axis is log_2_ fold change and Y axis is -Log_10_ of P value for B) Activated, C) Quiescent FOLR2^high^ and D) Quiescent FOLR^low^ Antigen Presenting IMs. Select genes with significant differential gene expression are labeled. E) Pie charts expressing the fraction of cells from each of the 5 IM clusters, split by smoking status of donor and lung structure of origin. F-H) Heat map of the percentages of IMs within each major hierarchal designation of Monocyte-like vs Antigen Presenting, Quiescent vs Inflammatory and FOLR2^low^ vs FOLR2^high^ split by smoking status [nonsmoker (NS) and smokers (S)] and structure-of-origin.

Next, we parsed apart IM clusters based on both structure-of-origin and smoking status to assess for differences in accumulation patterns, Figure 6E. There was a significant accumulation of Inflammatory IMs relatively to Quiescent IMs in the airways of smokers (66% [SD 27%]) versus nonsmokers (36% [SD 7%]), Figure 6F. Conversely, Quiescent IMs were more abundant than Inflammatory IMs in the alveoli of smokers (83% [SD 5%]) compared with nonsmokers (56% (SD 31%]). There were nonsignificant trends towards the accumulation of monocyte like IMs in the airways of smokers (56%) vs nonsmokers (38%) and FOLR2^Low^ IMs in the walls of vessels of smokers (43%) vs nonsmokers (27%). We lacked statical power to perform pairwise comparisons of transcriptional profiles to the level of IM subset, per structure-of-origin, per smoking status.

### Airspace Macrophages Exist in Conserved Transcriptional States

Next, we sought to define the spatial heterogeneity of transcriptionally defined airspace macrophage (AM) subsets within our RNA sequencing dataset. As a first step, we subclustered macrophages from all anatomical structures that were identified as AMs based on the expression of quintessential markers such as SPN and FABP4. This analysis identified five AM clusters, Figure 7A. Select, highly expressed genes characteristic of each population were identified, Figure 7B. Pathways analysis of the conserved markers for each AM population, Figure S5, suggest AM subsets aligned with previously identified populations. For example, AM1 cells showed enrichment in antigen processing and presentation pathways, while AM2 cells displayed elevated inflammatory responses and cytokine and chemokine expression. AM3 cells exhibited increased transcriptional activity, responsiveness to chemokines and cytokines, and negative regulation of apoptosis. AM4 cells were enriched for metallothionein-mediated metal processing, while AM5 cells exhibited a gene signature suggestive of interferon-responsiveness.

**Figure 7.**
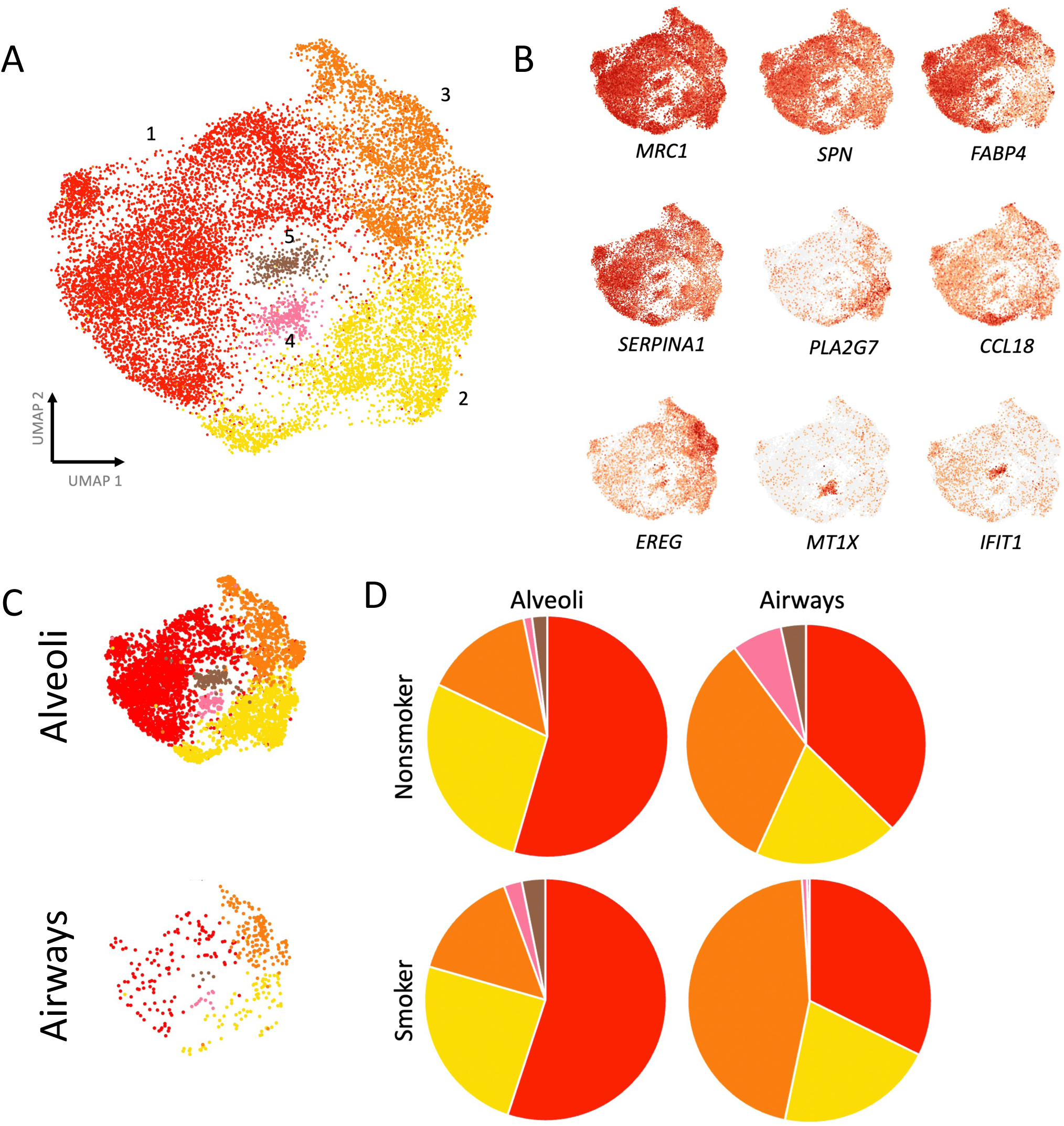
Airspace Macrophages (AMs) in Human lung tissue. UMAP projection of 5 AM populations defined by hierarchal clustering using the Seurat algorithm. B) Selected genes illustrating differential gene expression by select macrophage clusters. Colors are on a gradient from low (grey) to high (red) gene expression. C) UMAP of AMs split by compart-of-origin from Alveoli (top) or airways (bottom). D) Fractional accumulation of each AMs subset by donor smoking status (top/bottom) and structure of origin (left/right).

### Spatial Heterogeneity of Airspace Macrophages

To define the spatial heterogeneity of AMs transcriptionally, we analyzed AM subsets based on structure-of-origin hashtag to identify AMs arising from the alveoli or airways, Figure 7C. While all AM clusters were detectable in the alveolus and airways, AM1 cells were enriched in the alveolus, accounting for 55% of AMs. Interestingly, a different accumulation pattern emerged in the airways, where AM3 cells were most abundant, comprising 42% of AMs. Pathways analysis of AM1 suggests several antigen process and presentation pathways are highly enriched, in comparison to AM3 in which inflammatory response, cytokine signaling, and granulocyte recruitment response predominate. This suggests the general landscape is skewed to enhanced inflammation in the airways, in comparison to the alveoli. Due to the skew in AM composition between compartments, we did not have sufficient sample size to robustly compare gene expression between compartments within each AM subcluster

### Cigarette Smoke Results in Subtle Programming Changes to Airspace Macrophages

We compared the spatial distribution and programming of AMs from smokers versus nonsmokers to identify cigarette smoke-induced alterations. The fractional distribution of AM subpopulations from alveolar tissue was highly conserved amongst smokers and nonsmokers, Figure 7D. However, in the airways of cigarette smokers, we report a significantly increased fraction of AM3 (46% of AMs) compared to nonsmokers (33% of AMs). This increase in AM3 was accompanied by a corresponding reduction in the fraction of all other AM clusters in smokers, but particularly AM4, AM5 and AM6, Figure 7D.

Next, we performed pairwise comparisons of the transcriptional profiles of all AMs from smokers or nonsmokers. Surprisingly, only 7 significant DEGs were observed: in AMs from smokers, *MTMR11, KBTBD11, CD101, LTB4R*, and *ABCA5* were increased while *SCN9A* and *LINCO1914* were decreased. Next, we performed pairwise comparisons of the transcriptional profiles within each AM cluster between smokers vs nonsmokers, revealing a handful of DEGs within each. For example, 13 genes were up- and 6 genes were down-regulated in AM1 from smokers while 11 genes were up- and 13 genes were down-regulated in AM2 from smokers. The full list of genes altered by smoking in AMs is included in the supplemental materials.

## Discussion

The primary objective of this study was to categorize transcriptional profiles of macrophage populations within specific human lung anatomical structures. The resulting dataset provides a robust pool of human cells, enabling the identification of five interstitial macrophage (IM) and five airspace macrophage (AM) subsets distributed throughout, but with variable accumulation amongst different lung structures. Based on conserved gene expression signatures, subsets of IMs are classified within a novel three-tiered nomenclature: (1°) Antigen Presenting versus Monocyte-like, (2°) Inflammatory versus Quiescent, and (3°) *FOLR2*^high^ versus *FOLR2*^low^. Immunofluorescent microscopy confirms variable accumulation patterns of these IM subsets in lung anatomical structures. Finally, this study compared the distribution and expression patterns of lung macrophage subsets from nonsmokers and chronic cigarette smokers to detect subtle programming changes and highlight the enrichment of monocyte-like inflammatory IM and AM subsets in the airways of cigarette smokers.

Previous scRNA-seq studies conducted primarily in mice revealed that the expression of *Timd4, Lyve1*, and *Folr2* (T-L-F) broadly distinguish two subclasses of pulmonary interstitial macrophages (11, 15). In our human dataset, cells within the Quiescent and Inflammatory Antigen Presenting IM populations express *FOLR2* and exhibit similar transcription patterns to mouse *Cd163+/Folr2+* IMs (11) or *Timd4+/Lyve1+/Folr2+* IMs (17). *TIMD4* expression was absent in all human myeloid populations. *LYVE1* expression was observed in approximately half of the IMs which expressed FOLR2, Figure S6. Importantly, positive expression of the TLF genes FOLR2 and LYVE1 played only a minor role in distinguishing IM identity within our scRNA-seq dataset. Only 9 genes shared co-expression patterns with FOLR2 and LYVE1, suggesting a diminished importance (3° in our dataset) in defining IM identity. To emphasize this potential difference between human and mouse pulmonary IMs, we have avoided the TLF terminology to define human IMs in this dataset.

Lyve-1+ IMs, a subset of FOLR2^high^ IMs, have been reported to localize within vessel walls (8). Our study confirms but further elaborates upon this finding by highlighting the Inflammatory subset of FOLR2^high^ Antigen Presenting IMs, which express proinflammatory cytokines including CCL4, accumulates in the adventitia of the arterial wall. Many of those cells also express LYVE1 (Figure S6). Limited LYVE1 expression was detected outside of the arteries. CD169 (SIGLEC1) expression has been reported to mark a population of immune cells residing near nerves in murine lungs (nerve-airway macrophages, NAM (38)). Our scRNA-seq findings suggest that a significant portion of Antigen Presenting IMs, regardless of structure, express SIGLEC1. In fact, SIGLEC1 expression was more common amongst IMs which originated from the walls of arteries than airways (Figure S6).

We define a pulmonary Inflammatory IM subset based on a characteristic pattern of increased cytokine expression. Madissoon et. al. reported a population of human lungs IMs expressing CCL4 (21), and Li et. al. report that murine pulmonary IMs can be divided into numerous subsets based on cytokine expression profiles (18). Notably, our human IM populations exhibit high expression of CCL3, CCL4, CXCL2, and CXCL3, which aligns with the murine IMck2-4 subsets (18). However, we observed low expression of cytokines characteristic of several other murine IM subsets, such as IMck1 (CCL7, CXCL14), IMck5 (CCL8), IMck6 (Ccl6 and Ccl9 are exclusively expressed in mice), IMck7 (CXCL9, CXCL10), IMck8 (CXCL13), and IMck9 (CCL24) subtypes. Some of these populations may be absent due to a lack of acute inflammation in lung tissues analyzed in our study. Nevertheless, some differences may reflect biological differences between mice and humans. In support of this notion, we can identify fractions of cells consistent with each of our IM subsets, including Antigen Presenting, Monocyte-like, inflammatory and FOLR2^high^, in datasets published online (21, 36, 39).

Macrophage M1 and M2 polarization states have commonly been studied in vitro (40), but macrophages likely exist in more heterogeneous states in vivo (41, 42). The activated IM subsets in our dataset share some similarities with M1-polarized macrophages, including upregulation of IL1B. But some characteristic M1 markers, including iNOS, were not highly expressed in any human IM subsets. Our dataset lacks a population of IMs that clearly resemble M2 polarization. In our dataset, signatures of inflammatory transcriptional programming are present within subsets of both Antigen Presenting and Monocyte-like IMs. An explanation for this might be that the inflammatory state exists as a polarization option available to either subset of pulmonary IMs. However, it is intriguing that the inflammatory subset of Antigen Presenting IMs was conserved within the arterial wall across all donors analyzed by scRNA-seq and tissue histology. Such conservation of programming in this niche argues against the Inflammatory Antigen Presenting IM population existing as a transient polarization in response to, for example, an external stimulus. Further, the corroboration of Inflammatory IMs within the arterial walls by tissue histology argues against an alternative hypothesis of the inflammatory transcriptional state observed by scRNA-seq having been induced by warm tissue digestion or other cellular manipulation prior to sequencing (43). Future studies will investigate the significance of conserved Inflammatory lung IM subpopulations.

We hypothesized that human lung IM composition would be heavily influenced by anatomical structure-of-origin. This proved partially true. In keeping with recent observations in mouse tissues, we visualized accumulation of Frβ+ IMs throughout the bronchovascular bundles (8, 11, 16). However, none of the 5 IM subsets reported herein were restricted to a single structure. Rather, each IM subset was heterogeneouslydistributed throughout all lung structures. Interestingly, our microscopy findings suggest that the composition of IM subsets is determined on a more granular level than lung anatomical structure and is heavily influence by specific tissue layer niche. For instance, Frβ-negative IMs accumulate within the airway subepithelial lamina propria, while Frβ-positive IMs accumulate within the airway adventitia. Future studies should investigate which factors in the lung microanatomic niche drive IM heterogeneity.

Many aspects of the spatial distribution of macrophage subsets detected by scRNAseq or lung tissue histology are in general alignment. Both modalities suggest most IMs in the alveolus are Monocyte-like, Quiescent and FOLR2^low^ while IMs in the airways and vessels are more likely to be Antigen Presenting, Inflammatory and FOLR2^high^. However, in general, microscopy detected a higher percentage of CCL4+ Inflammatory IM and FOLR2^high^ IMs than predicted by scRNAseq. The divergence between scRNA-seq clusters and IF-based quantification likely reflects methodological and biological nuances. While scRNA-seq identifies macrophage subsets through combinatorial gene expression patterns, individual marker genes may exhibit broader protein expression across cell states or transient activation phases undetected by transcriptional thresholds (44). Likewise, technical factors, including antibody specificity and protein stability, combined with spatial microenvironmental influences on protein localization, may amplify IF signals in cells with low or sporadic transcript levels (45, 46). This underscores the importance of validating cluster-specific markers through orthogonal methods.

scRNA-seq analysis revealed that the fractional composition of pleural IMs detected did not significantly differ from alveolar tissue. However, image analysis suggests that CCL4-/Frβ- IMs accumulate within the alveolus but not the pleura. In addition to the limitations discussed above, we suspect that the detection of CCL4-/FOLR2^low^ IMs in pleural samples by scRNA-seq was due to contamination by alveolar cells. We acknowledge a limitation of this study: despite our best efforts to isolate individual tissue structures, cross-structure contamination likely occurred. We suspect that alveolar tissue, which directly abutted adjacent tissue structures including the pleura and bronchovascular bundles, was incompletely removed and contaminated other structures to some extent. Thus, tissue histology is likely a more accurate reflection of pleural macrophage composition than scRNA-seq in this study.

We report the accumulation of spatially distinct AM subsets, antigen-presenting AMs in the alveoli and monocyte-like AMs in the airway, which highlights the importance of understanding lung immune zonation. The heightened chemokine responsiveness in airway AMs suggests these cells are primed for rapid recruitment, potentially serving as sentinels at the bronchial-alveolar interface. Conversely, the antigen-processing specialization of alveolar AMs aligns with their role in surveilling pathogens while minimizing collateral inflammation. This spatial division of labor fits prior observations that airway-centric diseases such as COPD involve excessive monocyte recruitment (47), while alveolar pathologies such as pneumonia engage resident AMs’ phagocytic and regulatory functions (48).

Our identification of the accumulation of AMs and IMs with upregulated inflammatory signaling patterns in smokers’ airways align remarkably well with recent single-cell transcriptomic analyses demonstrating that cigarette smoke exposure drives recruitment of monocyte-derived alveolar macrophages with heightened inflammatory gene expression signatures (47). The gene expression patterns we observed suggest these cells are indeed recruited rather than resident macrophages, consistent with the emerging paradigm that monocyte-like AMs, rather than resident AM, are the predominant inflammatory drivers in smokers’ lungs. Our studies expand upon prior studies by highlighting that this accumulation occurs within the airway, without evidence of accumulation more diffusely throughout the alveoli. We will explore the implications of this finding in future studies.

While we observe changes in anatomical structural accumulation, we were surprised to find more modest programming differences in macrophages derived from chronic cigarette smokers compared with nonsmokers. This was surprising as metallothionein-expressing AMs have been observed to be increased in lungs of COPD patients (36). Our data suggest that the influence of cigarette smoke on macrophage programming may be more subtle than previously reported (49). However, an alternate explanation may be that cigarette smoke exposure results in relatively short-lived macrophage programming changes. We acknowledge that all tissue in this study was sourced from donors who were maintained on a ventilator, and obviously not exposed to cigarette smoke, for several days prior to organ donation. This multi-day delay between smoke exposure and our analyses may have provided adequate time for macrophages to lose programing changes induced by cigarette smoke. While the number of DEGs distinguishing IMs from smokers vs nonsmokers were limited, they suggest impairment in pathways crucial for host defense, including the response to interferon. While the roles of IMs in host defense are less extensively studied compared to their AMs counterparts (50), we hypothesize that smoke exposure impairs IMs’ ability to respond to pathogens, potentially contributing to the increased susceptibility to infection observed in chronic smokers (51–53).

In summary, we report the spatial distribution of human macrophages amongst major lung structures. We report five IM subsets detected within alveolar, pleural, airway or arterial walls and five AM subsets distributed throughout the alveoli and airway lumens. Human lung IM subsets are classified within a three-tiered system: (1°) Antigen Presenting vs Monocyte-like, (2°) Inflammatory vs Quiescent, and (3°) FOLR2^high^vs FOLR2^low^. While all IM subsets were observed in all lung substructures, the enrichment of each subset, and therefore the balance of IMs with characteristics of each classification tier, varied across anatomic regions. Monocyte-like IMs accumulate in alveolar parenchyma while Inflammatory Antigen Presenting IMs predominate in the bronchovascular bundles. A subset of Antigen Presenting AMs predominates in the alveoli while a more inflammatory monocyte-like population of AM is most prevalent in the airways. Surprisingly few programming differences were detected between macrophages sourced from chronic smokers vs nonsmokers but there was accumulation of inflammatory IM and AM subsets in the airways of smokers. By identifying the heterogeneous accumulation of macrophage subsets in major lung anatomical structures, our work informs future studies of the relationship between lung structure and the function of cells within while providing context for perturbations to these subsets in the diseased lung.

## Contributions

Conception and design- P.S.H, C.M.M, K.J.M, P.M.H, W.J.J.; Analysis and interpretation of data- P.S.H, E.W., K.H.L, C.M.M, W.J.J; Statistical methods and analysis- E.W., B.S., S.J., C.M.M.; Acquisition of data- P.S.H, K.H.L, J.D., Drafting and revising manuscript- P.S.H, E.W., K.H.L., K.J.M, C.M.M., W.J.J.

---

**Figure S1.**
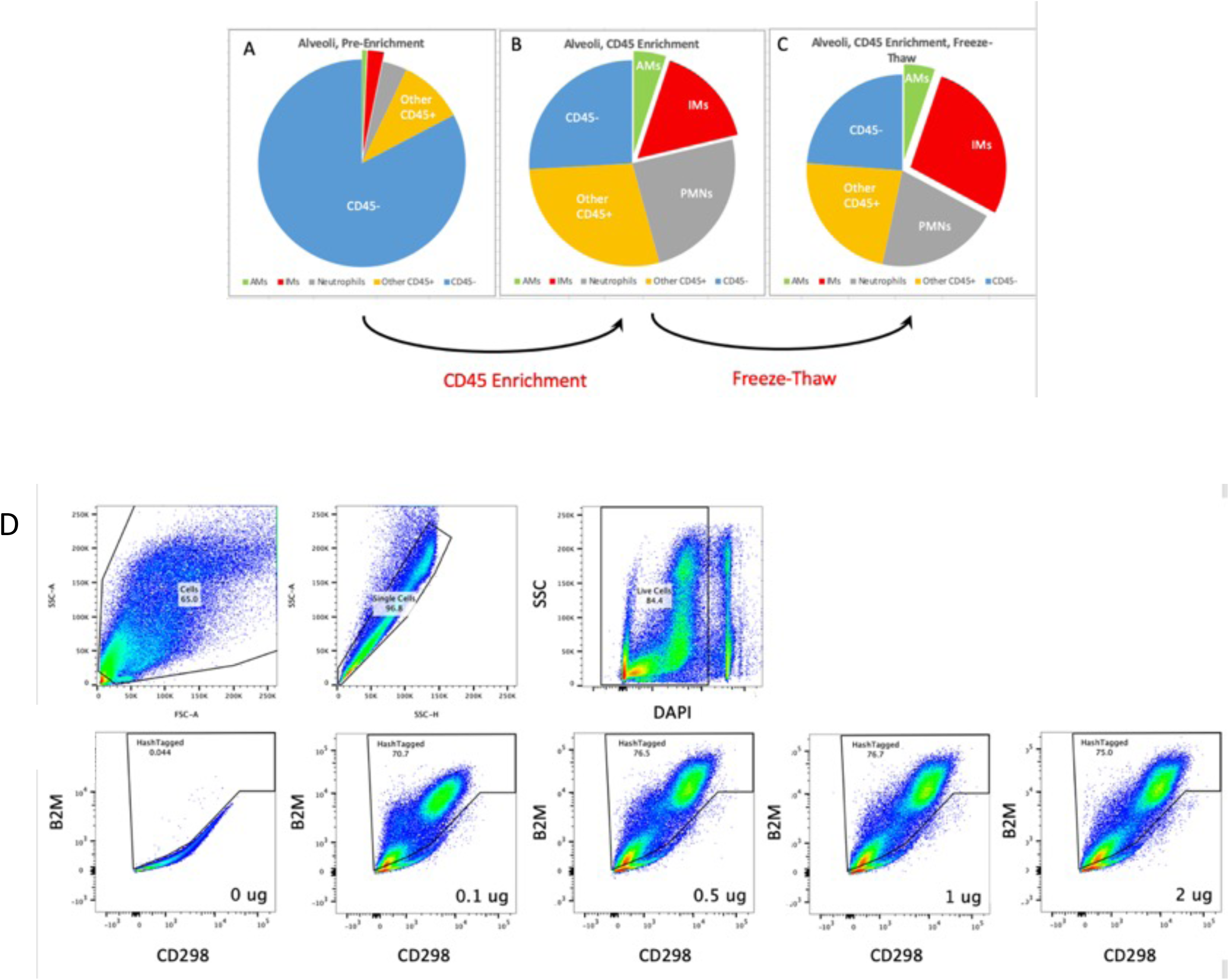
Validation of CD45 enrichment and cell hashing. A) Flow cytometry analysis of the fraction of live cells detected in human alveolar tissue after A) digest, followed by B) CD45- enrichment via magnetic bead separation, followed by C) freeze-thaw cycle. D) Flow cytometry of human digest cells labeled with anti-B2M and anti-CD298 antibodies analogous to the clones used in scRNA-seq experiments. Even low concentrations of antibodies (0.1 μg) resulted in labeling of the majority of cells.

**Figure S2.**
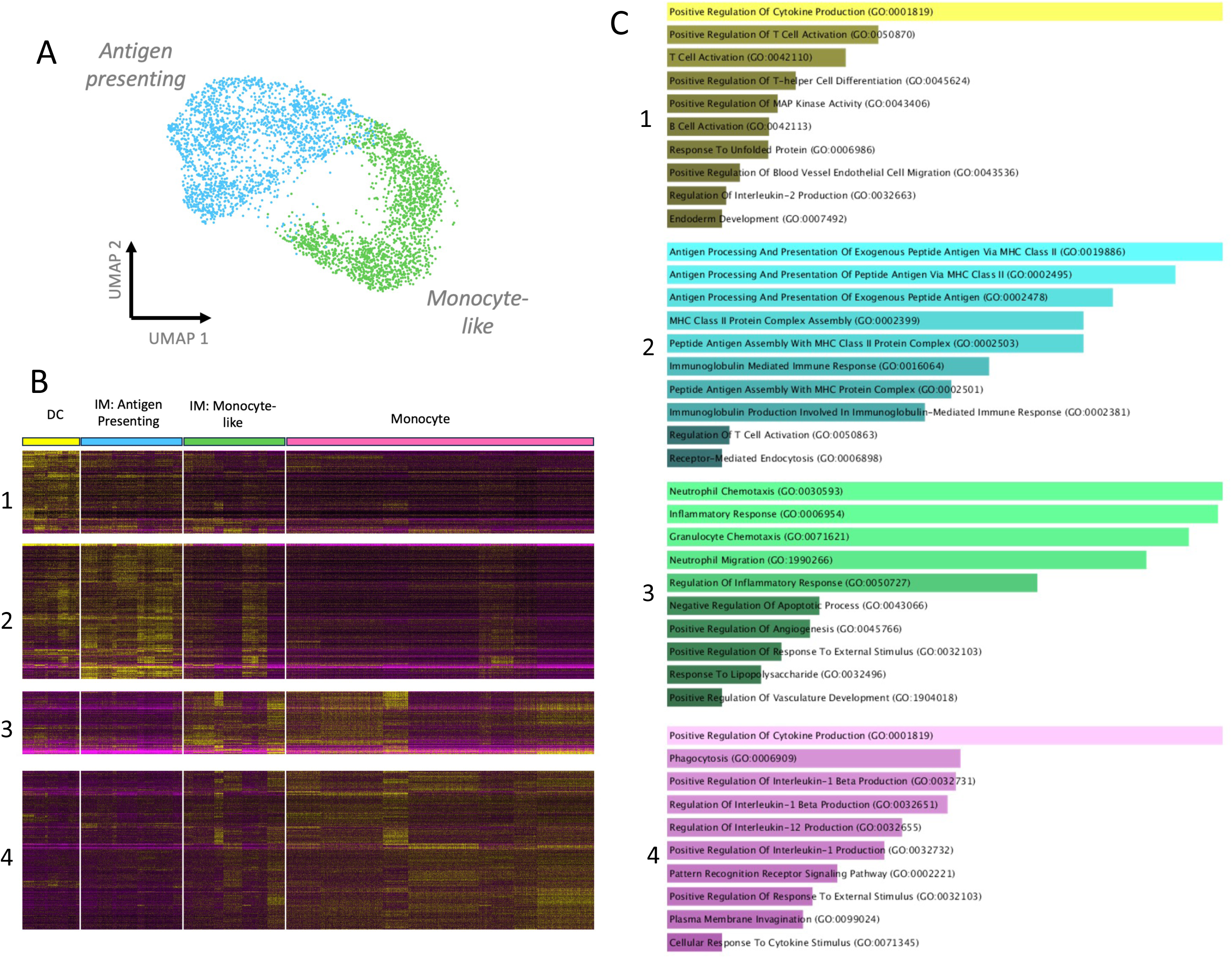
Two major populations of lung interstitial macrophages (IMs). A) UMAP projection of IMs highlighting two major populations: Antigen presenting (blue) and Monocyte-like (green) IMs. B) Heat map projection of all DEGs which emerged from all pairwise comparisons between dendritic cells (DC) antigen presenting IMs, Monocyte-like IMs and monocytes. C) Gene Ontology (GO) Pathways analysis of differentially expressed genes (DEGs) signatures from aforementioned heatmap highlighting the dominant pathways of each cell type: 1) dendritic cells (yellow), 2) Antigen Presenting IMs (blue), 3) Monocyte-like IMs (green), 4) Monocytes (purple).

**Figure S3.**
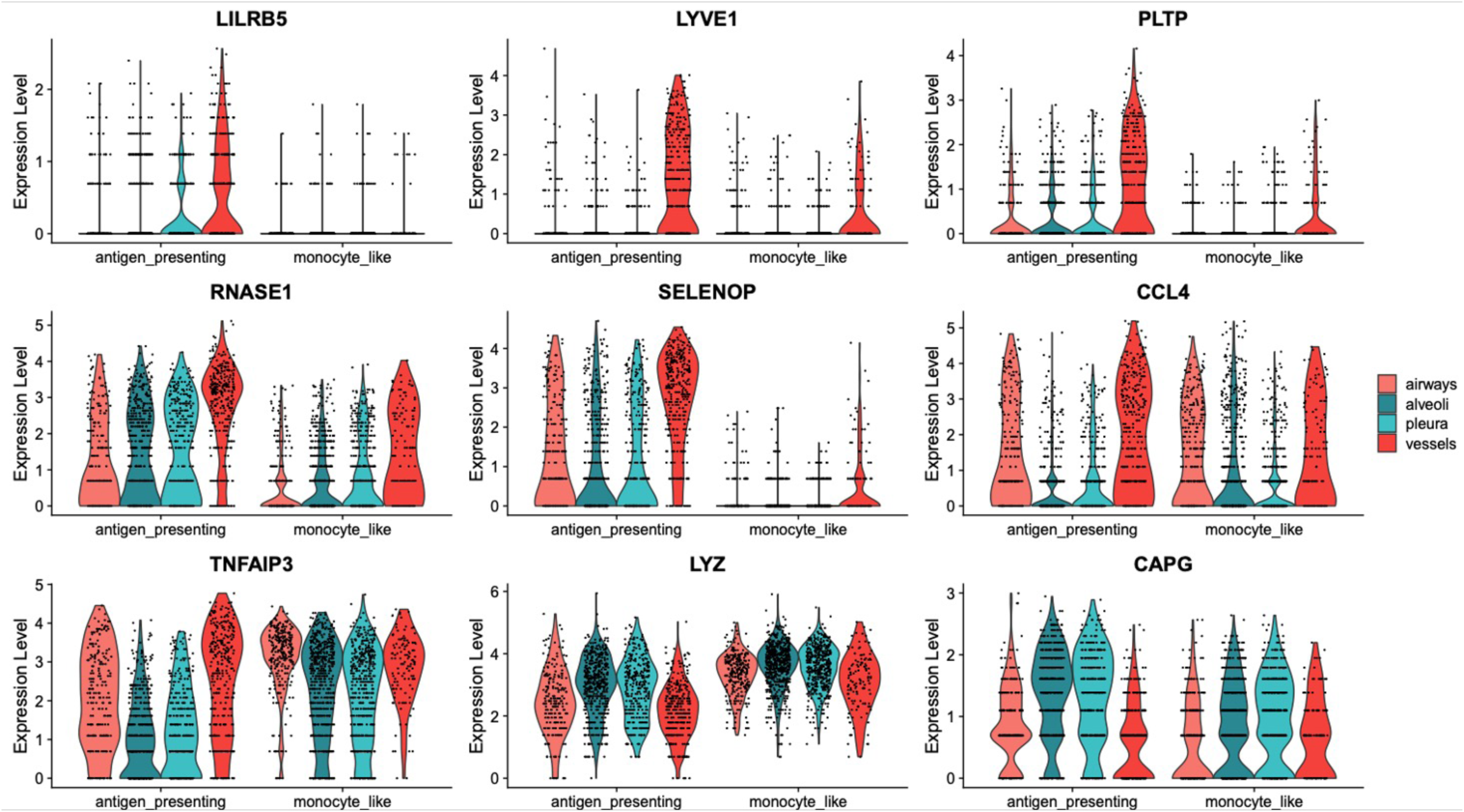
Violin plots of DEGs identified as distinguishing Antigen Presenting from Monocyte-like IM subsets based on structure of origin.

**Figure S4.**
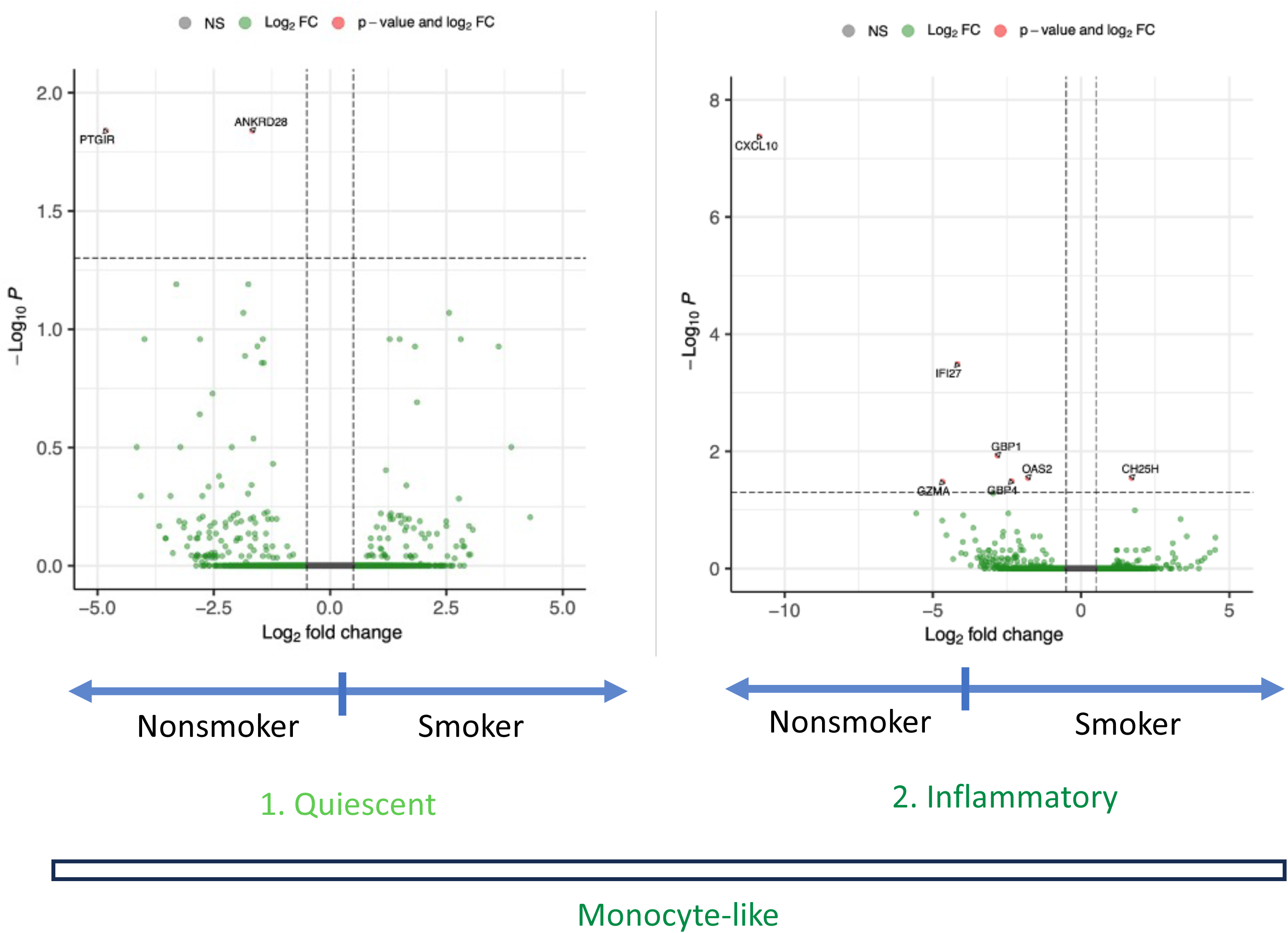
Volcano plots of all genes expressed in Monocyte-like IMs in nonsmoker vs smoking donors. Very few significant DEGs are identified in Left) Quiescent Monocyte-like IMs and Right) Inflammatory monocyte-like IMs.

**Figure S5.**
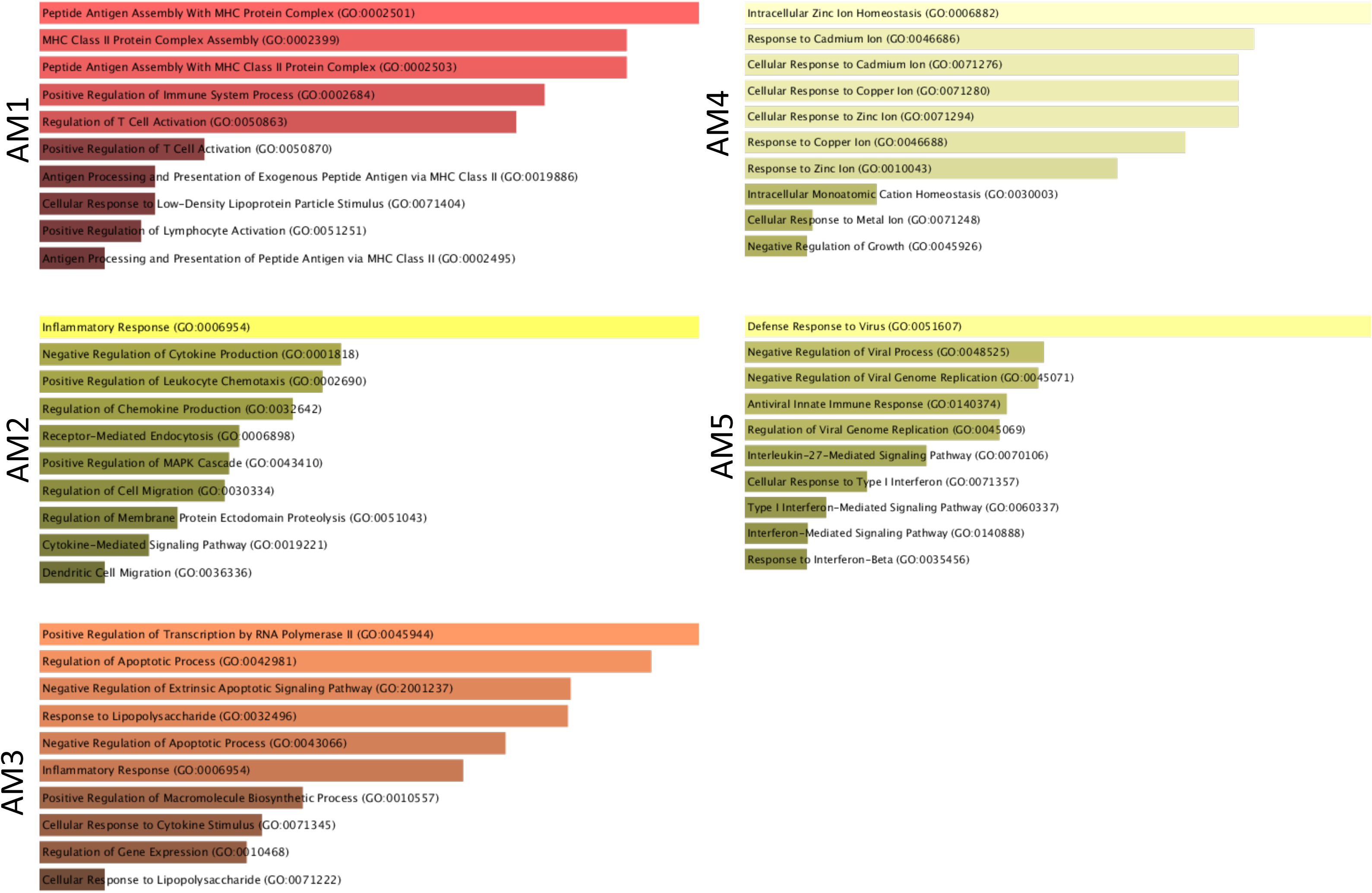
Gene Ontology (GO) Pathways analysis performed on all statistically significant (P<0.05) DEGs identified via all possibly pairwise comparisons of each airspace macrophage (AM) cluster. Highly enriched pathways shown for each AM cluster 1-5.

**Figure S6.**
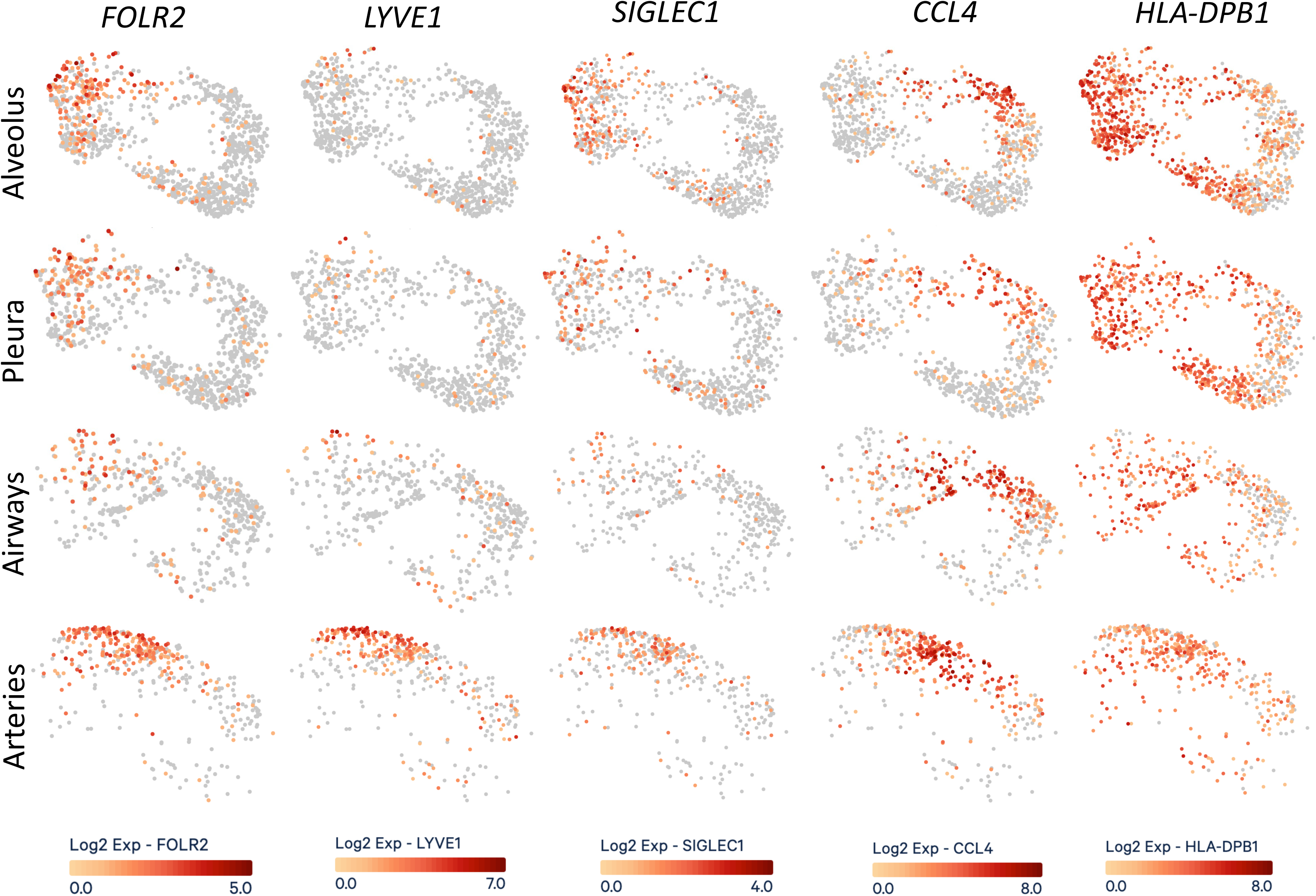
UMAP projection of all IMs, divided by structure of origin (vertically) and Log2 fold expression of *FOLR2, LYVE1, SIGLEC1* (CD169), *CCL4*, and *HLA-DPB1* (horizontally). Cells are colored on a gradient from grey (no gene expression) to red (high gene expression).

Table S1. Antibodies used for human flow cytometry (FC) and immunofluorescent (IF) histology.

## References

1. Hume PS, Gibbings SL, Jakubzick C V., Tuder RM, Curran-Everett D, Henson PM, et al. Localization of Macrophages in the Human Lung via Design-based Stereology. Am J Respir Crit Care Med 2020;201:1209–1217.

2. Amit I, Winter DR, Jung S. The role of the local environment and epigenetics in shaping macrophage identity and their effect on tissue homeostasis. Nat Immunol 2016;17:18– 25.

3. Ginhoux F, Jung S. Monocytes and macrophages: Developmental pathways and tissue homeostasis. Nat Rev Immunol 2014;14:392–404.

4. Murray PJ, Wynn TA. Protective and pathogenic functions of macrophage subsets. Nat Rev Immunol 2011;11:723–737.

5. Wynn TA, Vannella KM. Macrophages in Tissue Repair, Regeneration, and Fibrosis. Immunity 2016;44:450–462.

6. Wang Y, Xu J, Meng Y, Adcock IM, Yao X. Role of inflammatory cells in airway remodeling in COPD. International Journal of COPD 2018;13:3341–3348.

7. Sabatel C, Radermecker C, Fievez L, Paulissen G, Chakarov S, Fernandes C, et al. Exposure to Bacterial CpG DNA Protects from Airway Allergic Inflammation by Expanding Regulatory Lung Interstitial Macrophages. Immunity 2017;46:457–473.

8. Chakarov S, Lim HY, Tan L, Lim SY, See P, Lum J, et al. Two distinct interstitial macrophage populations coexist across tissues in specific subtissular niches. Science (1979) 2019;363:.

9. Pullamsetti SS, Savai R. Macrophage regulation during vascular remodeling: Implications for pulmonary hypertension therapy. Am J Respir Cell Mol Biol 2017;56:556–558.

10. Gosselin D, Link VM, Romanoski CE, Fonseca GJ, Eichenfield DZ, Spann NJ, et al. Environment drives selection and function of enhancers controlling tissue-specific macrophage identities. Cell 2014;159:1327–1340.

11. Moore PK, Anderson KC, McManus SA, Tu T-H, King EM, Mould KJ, et al. Single-Cell RNA Sequencing Reveals Unique Monocyte-derived Interstitial Macrophage Subsets during Lipopolysaccharide-Induced Acute Lung Inflammation. Am J Physiol Lung Cell Mol Physiol 2023;doi:10.1152/ajplung.00223.2022.

12. Mould KJ, Jackson ND, Henson PM, Seibold M, Janssen WJ. Single cell RNA sequencing identifies unique inflammatory airspace macrophage subsets. JCI Insight 2019;4:.

13. Mould KJ, Moore CM, McManus SA, McCubbrey AL, McClendon JD, Griesmer CL, et al. Airspace macrophages and monocytes exist in transcriptionally distinct subsets in healthy adults. Am J Respir Crit Care Med 2021;203:946–956.

14. Li X, Kolling FW, Aridgides D, Mellinger D, Ashare A, Jakubzick C V. ScRNA-seq expression of IFI27 and APOC2 identifies four alveolar macrophage superclusters in healthy BALF. Life Sci Alliance 2022;5:1–17.

15. Li X, Mara AB, Musial S, Rawat K, King WT, Kolling FW, et al. Coordinated Chemokine Expression Defines Macrophage Subsets Across Tissues. 2023;1–48.at <10.1101/2023.05.12.540435>.

16. Gibbings SL, Thomas SM, Atif SM, McCubbrey AL, Desch AN, Danhorn T, et al. Three Unique Interstitial Macrophages in the Murine Lung at Steady State. Am J Respir Cell Mol Biol 2017;57:1–44.

17. Dick SA, Wong A, Hamidzada H, Nejat S, Nechanitzky R, Vohra S, et al. Three tissue resident macrophage subsets coexist across organs with conserved origins and life cycles. Sci Immunol 2022;7:eabf7777.

18. Li X, Mara AB, Musial SC, Kolling FW, Gibbings SL, Gerebtsov N, et al. Coordinated chemokine expression defines macrophage subsets across tissues. Nat Immunol 2024;206:.

19. Leach SM, Gibbings SL, Tewari AD, Atif SM, Vestal B, Danhorn T, et al. Human and Mouse Transcriptome Profiling Identifies Cross-Species Homology in Pulmonary and Lymph Node Mononuclear Phagocytes. Cell Rep 2020;33:108337.

20. Desch AN, Gibbings SL, Goyal R, Kolde R, Bednarek J, Bruno T, et al. Flow Cytometric Analysis of Mononuclear Phagocytes in Nondiseased Human Lung and Lung-Draining Lymph Nodes. Am J Respir Crit Care Med 2016;193:614–26.

21. Madissoon E, Oliver AJ, Kleshchevnikov V, Wilbrey-Clark A, Polanski K, Richoz N, et al. A spatially resolved atlas of the human lung characterizes a gland-associated immune niche. Nat Genet 2023;55:66–77.

22. Gibbings SL, Jakubzick C V. Isolation and Characterization of Mononuclear Phagocytes in the Mouse Lung and Lymph Nodes. Methods Mol Biol 2018;1809:33–44.

23. Atif SM, Gibbings SL, Jakubzick C V. Isolation and Identification of Interstitial Macrophages from the Lungs Using Different Digestion Enzymes and Staining Strategies. Methods Mol Biol 2018;1784:69–76.

24. Cohen MC. LEUKOCYTE AND STROMAL CELL MOLECULES. Shock 2007;28:498.

25. Zheng GXY, Terry JM, Belgrader P, Ryvkin P, Bent ZW, Wilson R, et al. Massively parallel digital transcriptional profiling of single cells. Nat Commun 2017;8:.

26. Hao Y, Hao S, Andersen-Nissen E, Mauck WM, Zheng S, Butler A, et al. Integrated analysis of multimodal single-cell data. Cell 2021;184:3573–3587.e29.

27. 27. Peripheral blood mononuclear cells (PBMCs from a healthy donor. At <https://cf.10xgenomics.com/samples/cell-vdj/6.1.0/10k_PBMC_5pv2_nextgem_Chromium_Controller_10k_PBMC_5pv2_nextgem_Chromium_Controller/10k_PBMC_5pv2_nextgem_Chromium_Controller_10k_PBMC_5pv2_nextgem_Chromium_Controller_web_summary.html>.

28. Murtagh F, Legendre P. Ward’s Hierarchical Agglomerative Clustering Method: Which Algorithms Implement Ward’s Criterion? J Classif 2014;31:274–295.

29. Lun ATL, Marioni JC. Overcoming confounding plate effects in differential expression analyses of single-cell RNA-seq data. Biostatistics 2017;18:451–464.

30. Crowell HL, Soneson C, Germain PL, Calini D, Collin L, Raposo C, et al. Muscat Detects Subpopulation-Specific State Transitions From Multi-Sample Multi-Condition Single-Cell Transcriptomics Data. Nat Commun 2020;11:1–12.

31. Robinson MD, McCarthy DJ, Smyth GK. edgeR: A Bioconductor package for differential expression analysis of digital gene expression data. Bioinformatics 2009;26:139–140.

32. Kuleshov M V., Jones MR, Rouillard AD, Fernandez NF, Duan Q, Wang Z, et al. Enrichr: a comprehensive gene set enrichment analysis web server 2016 update. Nucleic Acids Res 2016;44:W90–W97.

33. Hao Y, Hao S, Andersen-Nissen E, Mauck WM, Zheng S, Butler A, et al. Integrated analysis of multimodal single-cell data. Cell 2021;184:3573–3587.e29.

34. Vieira Braga FA, Kar G, Berg M, Carpaij OA, Polanski K, Simon LM, et al. A cellular census of human lungs identifies novel cell states in health and in asthma. Nat Med 2019;25:1153–1163.

35. Mulder K, Patel AA, Kong WT, Piot C, Halitzki E, Dunsmore G, et al. Cross-tissue single-cell landscape of human monocytes and macrophages in health and disease. Immunity 2021;1–18.doi:10.1016/j.immuni.2021.07.007.

36. Sauler M, McDonough JE, Adams TS, Kothapalli N, Barnthaler T, Werder RB, et al. Characterization of the COPD alveolar niche using single-cell RNA sequencing. Nat Commun 2022;13:494.

37. Travaglini KJ, Nabhan AN, Penland L, Sinha R, Gillich A, Sit R V., et al. A molecular cell atlas of the human lung from single-cell RNA sequencing. Nature 2020;587:619–625.

38. Ural BB, Yeung ST, Damani-Yokota P, Devlin JC, de Vries M, Vera-Licona P, et al. Identification of a nerve-associated, lung-resident interstitial macrophage subset with distinct localization and immunoregulatory properties. Sci Immunol 2020;5:.

39. Adams TS, Schupp JC, Poli S, Ayaub EA, Neumark N, Ahangari F, et al. Single-cell RNA-seq reveals ectopic and aberrant lung-resident cell populations in idiopathic pulmonary fibrosis. Sci Adv 2020;6:.

40. Murray PJ, Allen JE, Biswas SK, Fisher EA, Gilroy DW, Goerdt S, et al. Macrophage Activation and Polarization: Nomenclature and Experimental Guidelines. Immunity 2014;41:14–20.

41. Strizova Z, Benesova I, Bartolini R, Novysedlak R, Cecrdlova E, Foley LK, et al. M1/M2 macrophages and their overlaps - myth or reality? Clin Sci 2023;137:1067–1093.

42. Mould KJ, Barthel L, Mohning MP, Thomas SM, McCubbrey AL, Danhorn T, et al. Cell Origin Dictates Programming of Resident Versus Recruited Macrophages During Acute Lung Injury. Am J Respir Cell Mol Biol 2017;57:rcmb.2017-0061OC.

43. Denisenko E, Guo BB, Jones M, Hou R, De Kock L, Lassmann T, et al. Systematic assessment of tissue dissociation and storage biases in single-cell and single-nucleus RNA-seq workflows. Genome Biol 2020;21:1–25.

44. Squair JW, Gautier M, Kathe C, Anderson MA, James ND, Hutson TH, et al. Confronting false discoveries in single-cell differential expression. Nat Commun 2021;12:.

45. Sorokin M, Ignatev K, Poddubskaya E, Vladimirova U, Gaifullin N, Lantsov D, et al. RNA sequencing in comparison to immunohistochemistry for measuring cancer biomarkers in breast cancer and lung cancer specimens. Biomedicines 2020;8:.

46. Peng H, Wu X, Liu S, He M, Xie C, Zhong R, et al. Multiplex immunofluorescence and single-cell transcriptomic profiling reveal the spatial cell interaction networks in the non-small cell lung cancer microenvironment. Clin Transl Med 2023;13:.

47. Wohnhaas CT, Baßler K, Watson CK, Shen Y, Leparc GG, Tilp C, et al. Monocyte-derived alveolar macrophages are key drivers of smoke-induced lung inflammation and tissue remodeling. Front Immunol 2024;15:1–24.

48. Rubins JB. Alveolar macrophages: wielding the double-edged sword of inflammation. Am J Respir Crit Care Med 2003;167:103–4.

49. Liegeois M, Bai Q, Fievez L, Pirottin D, Legrand C, Guiot J, et al. Airway Macrophages Encompass Transcriptionally and Functionally Distinct Subsets Altered by Smoking. Am J Respir Cell Mol Biol 2022;67:241–252.

50. Lugg ST, Scott A, Parekh D, Naidu B, Thickett DR. Cigarette smoke exposure and alveolar macrophages: Mechanisms for lung disease. Thorax 2022;77:94–101.

51. Vanker A, Gie RP, Zar HJ. The association between environmental tobacco smoke exposure and childhood respiratory disease: a review. Expert Rev Respir Med 2017;11:661–673.

52. Ghio AJ. Particle exposures and infections. Infection 2014;42:459–467.

53. Bowsher R, Marczylo TH, Gooch K, Bailey A, Wright MD, Marczylo EL. Smoking and vaping alter genes related to mechanisms of SARS-CoV-2 susceptibility and severity. European Respiratory Journal 2024;2400133.doi:10.1183/13993003.00133-2024.

